# Integrating fossils samples with heterogeneous diversification rates: a combined Multi-Type Fossilized Birth-Death model

**DOI:** 10.1101/2025.02.03.636219

**Authors:** Joëlle Barido-Sottani, Hélène Morlon

## Abstract

Birth-death models are widely used to describe the diversification process which leads to the observed species and phylogenies. When integrated into Bayesian phylogenetic inference, birth-death models allow the joint inference of the phylogeny and the diversification parameters from molecular information. Two major classes of extensions of the birth-death process are considered in this article. The first extends the phylogenetic tree to include fossil samples alongside extant species, allowing the inference to integrate information about the past diversity. The second extension models diversification rates which can vary between lineages, and is thus able to infer patterns of variation in speciation or extinction rates. In this work, we combine these two types of extension into a multi-type fossilized birth-death (MTFBD) process, which can perform the joint inference of a phylogeny including extinct and extant samples, and lineage-specific diversification and fossilization rates in a Bayesian framework. The MTFBD model is implemented as part of the phylogenetic inference framework BEAST2. Using simulated and empirical datasets, we demonstrate the performance and accuracy of the new model compared to a model with rate heterogeneity but using only extant samples, and compared to a model without rate variation including fossils. We demonstrate that including fossils improves the accuracy of the phylogeny and diversification rates, especially extinction rates, provided that the inference includes morphological information to accurately place the fossil samples. When this information is not available however, MTFBD estimates are strongly driven by the priors and are thus no better or even worse than estimates obtained only with extant samples. With informative fossil characters, the MTFBD model provides accurate phylogenies, and precisely characterizes how speciation, extinction and fossilization rates vary as diversification proceeds.

## 1 Introduction

Time-calibrated trees represent the evolutionary relationships between species or organisms as well as a timescale of these relationships, therefore providing a crucial basis for hypothesis-testing in the life and earth sciences. Phylogenetic inference uses molecular alignment data to estimate phylogenetic trees, and can be implemented in a Maximum Likelihood (ML) or Bayesian framework. One specificity of Bayesian phylogenetic inference is that it includes a model of the evolutionary process which gave rise to the phylogeny, also called a tree prior. In full phylogenetic inference, the tree prior and the phylogeny are estimated jointly from molecular data, using for instance the BEAST2 software framework (Bouckaert et al., 2019).

Tree priors are generally divided into two main classes, coalescent processes and birth-death processes. This second category assumes that the tree process is driven by two main evolutionary rates: the birth rate, which describes the rate at which new lineages arise in the phylogeny, and the death rate, which describes the rate at which lineages are removed from the phylogeny (MacPherson et al., 2021). Birth-death models are also widely applied directly to a previously estimated phylogeny to infer the underlying evolutionary process (Pennell and Harmon, 2013; Stadler, 2013; Morlon, 2014; Morlon et al., 2024).

Many different extensions have been developed for the basic birth-death model to offer a better representation of the actual diversification process (Morlon et al., 2024). One important category of extended birth-death models are heterogeneous models, which integrate variation in birth and death rates across lineages. Such variation is generally assumed to be widespread in empirical datasets, and many morphological and ecological traits have been proposed to contribute to this variability, such as body size (Inostroza-Michael et al., 2018) or environmental conditions (Letsch et al., 2018). Birth-death processes which have been developed to account for across-lineage variation in rates assume either that the evolutionary process can be divided into several discrete diversification regimes (within which lineages share birth and death rates, see Maddison et al. (2007); FitzJohn (2012); Vasconcelos et al. (2022); Rabosky et al. (2013); Kühnert et al. (2016); Höhna et al. (2019); Barido-Sottani et al. (2020b)), or that each lineage has its own birth and death rates, which vary cladogenetically (Maliet et al., 2019; Maliet and Morlon, 2022) or anagenetically (Quintero et al., 2024). We focus here on the first type of models, which can itself be divided in two main categories: one category of model assumes that there is a known trait, or combination of traits, fixed by the user, which drives rate variations, while the second category assumes that this information is not available. We focus here on the latter, which include the methods BAMM (Rabosky et al., 2013), LSBDS (Höhna et al., 2019), MSBD (Barido-Sottani et al., 2020b) and MiSSE (Vasconcelos et al., 2022). These methods differ by their implementations and assumptions, but share some common characteristics. In particular, they divide the evolutionary process into several discrete diversification regimes, each characterized by its own birth and death rates. Except for BAMM, these methods also include a transition process associated with one or several type change rate parameters. The total number of regimes can be estimated (BAMM, MSBD) or set to a fixed number chosen by the user (LSBDS, MiSSE). Overall, all methods allow users to estimate variations in diversification rates across a phylogeny, without needing to make assumptions on the underlying trait driving these patterns.

A second category of extensions deals with the inclusion of fossil samples into the phylogeny. Fossil evidence is a crucial source of information for dating precisely evolutionary events, including phylogenetic divergence times. The fossilised birth-death (FBD) process explicitly combines the lineage diversification and fossil recovery processes, providing an ideal model for inferring phylogenies from both extant species and fossil specimens (Stadler, 2010). Using this approach, fossils (with or without character data) are directly considered as part of the tree (Heath et al., 2014; Gavryushkina et al., 2014, 2017; Zhang et al., 2016). Since molecular sequences are very rarely available for extinct species, additional information is required to infer their position in the phylogeny. This information can be provided in two different forms, which can be used separately or in combination (Barido-Sottani et al., 2023). The first is topological constraints, which are constraints added by the user specifying that certain subsets of tips, extant or extinct, need to be monophyletic clades in the inferred phylogeny. These constraints are generally based on the taxonomy, with the constrained clades corresponding to genera and/or higher classifications. Topological constraints do not directly contribute to the likelihood calculation, but restrict the tree space which can be explored by the inference. The second approach, called a total-evidence analysis (Ronquist et al., 2016), uses a morphological data matrix with data for both fossil and extant specimens, in addition to the molecular alignment. The morphological matrix is added to the inference along with a morphological substitution model and a morphological clock model, and contributes to the phylogenetic likelihood. Previous work has shown that both approaches can infer accurate phylogenies under the FBD process (Barido-Sottani et al., 2023). Overall, the FBD process is a statistically coherent prior on divergence times, where the variance associated with node ages reflects the incompleteness of the fossil record, as well as uncertainty associated with the placement of fossil taxa in the phylogeny. Bayesian inference using the FBD process as a tree prior also allows for more reliable estimation of the diversification dynamics, particularly regarding extinctions. The FBD process is implemented in most major phylogenetic inference tools, including BEAST2 as the Sampled Ancestors (SA) package (Gavryushkina et al., 2014). An extension of the FBD model, the Occurrence Birth-Death (OBD) process (Andréoletti et al., 2022), has been developed to account for occurrences, i.e. fossil samples whose age is known but which cannot be placed in a phylogeny due to missing taxonomical or morphological information. However, this model has not been widely adopted due in part to its computational cost.

Although the FBD and MTBD processes have generally been developed and used separately, they are by no means incompatible. Indeed, the inclusion of fossil samples or across-lineage rate heterogeneity are more accurately considered as different features of the same very general birth-death model, rather than different models (MacPherson et al., 2021). Several implementations of rate-heterogeneous models have been extended to account for fossil samples, including fossil-BAMM (Mitchell et al., 2019) and the MiSSE process (Beaulieu and O’Meara, 2023). Results concerning the effect of including extinct species have been mixed, with one study finding improvement in the accuracy of extinction rate estimates (Mitchell et al., 2019), while another found that including fossil samples did not improve the accuracy of diversification rate estimates and could even be detrimental (Beaulieu and O’Meara, 2023). However, these implementations are limited by several factors. First, they can only be applied to fixed phylogenies. The phylogenetic position of fossil samples is rarely known exactly, even for well-studied groups such as the cetaceans (Marx et al., 2016), so assuming a fixed phylogeny is likely unrealistic for many empirical datasets. Second, both implementations only include heterogeneity in the birth and death rates, and assume that the fossilization process is homogeneous.

In this article, we present a full implementation of a combined MTFBD process, including fossilization rate heterogeneity, which can be used to co-estimate the phylogeny, the diversification and the fossilization process, either using topological constraints or a total-evidence approach. We use it to study how combined models accounting for both fossil samples and rate heterogeneity perform in a context where the phylogeny is not assumed to be known, but is estimated from molecular and morphological or taxonomical information. We demonstrate the performance and accuracy of this new model compared to either the original MTBD process (with rate heterogeneity but without fossils) or the original FBD process (with fossil samples but homogeneous), on both simulated and empirical datasets. Our MTFBD implementation is available as an extension of the BEAST2 package MSBD, and can thus be combined with all other functions and models already present in BEAST2.

## 2 Methods

### 2.1 Combined MTFBD model

The combined MTFBD model is based on the MTBD model described in Barido-Sottani et al. (2020b). The model contains a discrete number *n*\* of types, which correspond to distinct diversification regimes. Each type *i* is associated with its own birth rate λ_*i*_, its own death rate *µ*_*i*_, and specifically for the MTFBD model its own fossil sampling rate *ψ*_*i*_. We assume that death and fossil sampling events are completely independent, i.e. a lineage always continues after being sampled. This corresponds to a fixed removal probability of *r* = 0 in the general birth-death model presented by MacPherson et al. (2021). At each birth event, the type of the ancestral lineage is inherited by the two descendant lineages. Type change events can happen anywhere in the phylogeny, following a Poisson process with parameter the type change rate *γ*. At these type change events, a new type is selected uniformly at random among the *n*\* − 1 available. In particular, this means that the same type can be present in multiple subclades of the phylogeny. Note also that the total number of types *n*\* can be greater than the number of types which actually appear in the sampled tree.

The process starts at the root *T* = *t*_*origin*_ with a type selected uniformly at random, and continues until the present *T* = 0. Lineages which survive to the present are sampled with probability *ρ*. The final reconstructed tree is composed of all lineages which have at least one sampled descendant, extant or extinct. Fossil samples which have sampled descendants are called “sampled ancestors” (SAs), and by convention are represented as tips at the end of a branch of length 0. Fossil samples which do not have sampled descendants are called “fossil tips”. The type of a sampled fossil always matches with the type of the lineage from which it was sampled.

To calculate the likelihood, we define *p*_*i*_(*t*) the probability that a lineage in type *i* at time *t* will not leave any sampled descendants, and *q*_*i*_(*t*_*s*_, *t*_*e*_) the probability that a lineage in type *i* at time *t*_*s*_ will undergo no sampled events until time *t*_*e*_ (i.e. this lineage will be recorded as a single continuous edge in the final sampled tree). In the MTFBD process, these two quantities follow the master equations outlined here (see MacPherson et al. (2021) for a full derivation):

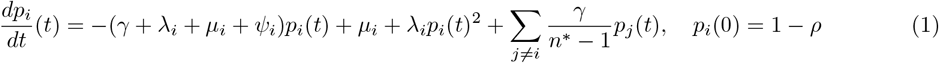

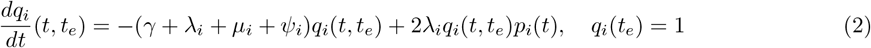

The full likelihood for the sampled phylogeny 𝒯 and the associated types 𝒮 given the parameters of the model *η* = (*λ, µ, γ, ψ, n*\*) is then

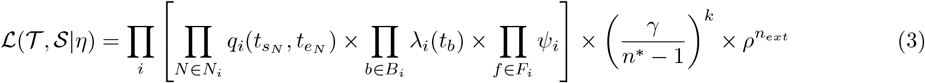

where *N*_*i*_ is the set of edges in type *i*, 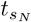 and 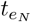 are respectively the start and end times of edge *N, B*_*i*_ is the set of birth events in type *i, F*_*i*_ is the set of fossil samples in type *i* and *n*_*ext*_ is the number of extant samples.

The set of master equations (1) and (2) do not have a close form solution. Some implementations of the MTBD model, such as the BEAST2 package BDMM (Kühnert et al., 2016), use numerical integration techniques to calculate the likelihood, however this process is computationally expensive and can lead to numerical instabilities in some parts of the parameter space. Instead, we make the assumption that type change events only appear in sampled parts of the tree. This assumption was also present in the original MTBD model described in Barido-Sottani et al. (2020b), and was found to have a limited impact on the results of the inference, while significantly improving the computational efficiency of the model. Following this assumption, equation (1) then becomes

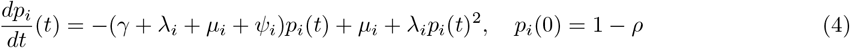

which can be solved analytically.

The closed-form solutions for *p*_*i*_ and *q*_*i*_ are then as follows:

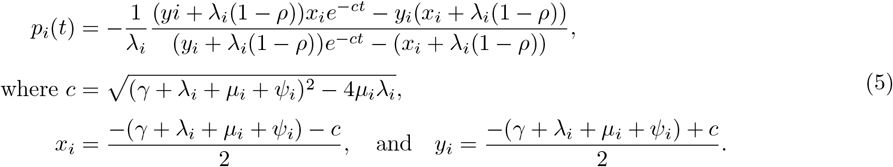

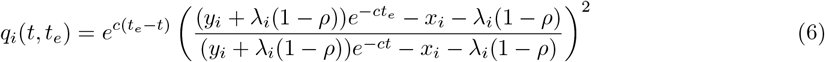

A more complete description of both models involved, as well as the derivation of the likelihood equations, can be found in Stadler (2010) for the FBD process and Barido-Sottani et al. (2020b) for the MTBD process. A complete derivation of the likelihood of a general birth-death-sampling process with types can be found in MacPherson et al. (2021).

### 2.2 Implementation of the MTFBD model

In addition to the changes to the likelihood described in the previous section, the implementation of the MTFBD model required several changes to the proposal algorithms used in the MCMC to propose new parameter states. In particular, sampled ancestors need to be handled differently than normal internal nodes, since they represent sampling events rather than birth events. We also implemented new proposals to sample the ages of the fossil tips, as well as estimate whether any fossil is a tip or a sampled ancestor. These proposals were based on existing proposals in the SA package, extended to account for the different types present in the MTBD model.

Similar to the original MTBD model, the MTFBD implementation outputs so-called “coloured” trees, where all edges and samples are assigned to a specific type. It also provides estimates for the total number of types present in the process, and birth, death and fossil sampling rates associated with each type. Following Barido-Sottani et al. (2019, 2020a), the model allows users to integrate fossil age uncertainty by providing age distributions on each fossil sample.

The extended model is implemented in the BEAST2 package MSBD, starting from version v2.0.3. The package can be installed directly through the BEAUti interface or from the Bitbucket repository https://bitbucket.org/bjoelle/msbd/. The combined model is fully integrated into the BEAUti graphical interface, allowing users to switch between the original model and the extended model with fossils based on their dataset. As demonstrated in the following simulation study, the MTFBD implementation can be easily combined with other components of the BEAST2 framework, for instance a morphological substitution model to perform total-evidence analyses.

### 2.3 Validation

We validated our new MTFBD implementation by using the simulation-based calibration procedure, introduced by Talts et al. (2018) and further described by Andréoletti et al. (2022). This procedure combines a simulation and an inference step, in order to ensure that the model used for the inference matches with the simulation model. In the simulation step, we simulated 1000 phylogenies with extant and extinct samples following the MTFBD model. For each replicate, the origin time, the number of types in the MTFBD model and the MTFBD rates were drawn from the same distributions. The full list of the distributions used can be found in Table S1. For all replicates, we then simulated sequence alignments for both extinct and extant samples, using a strict clock with clock rate *c* = 0.03 and a Jukes-Cantor substitution model. Note that we use these alignments purely for validating our implementation of the MTFBD model, and so we can use molecular sequences for the fossil samples rather than the morphological sequences which would be used in a real inference, as long as the simulation and inference models match perfectly. Finally, we used the alignments to perform an inference under our new MTFBD implementation. In the inference, the clock and substitution models were fixed to the simulation model (including a fixed clock rate), and the priors on the other parameters were set to the distributions used for sampling in the simulation step. We then checked whether the true simulated parameters fell into the inferred credible interval for each replicate. If both simulation and inference models match, we expect that *α*% of replicates will fall into the *α*% credible interval, for any value of *α*. As shown in Figure S1, all tested parameters follow this pattern, validating that our MTFBD implementation is correct.

### 2.4 Simulation study

#### 2.4.1 Simulating trees

Phylogenies were simulated under a forward birth-death process, using the generalized sampling approach (Hartmann et al., 2010) to obtain *n*_*tips*_ = 25 at the present. Fossil samples were then simulated using a Poisson process on the full phylogeny. The sampled phylogeny was obtained by discarding the unsampled lineages, i.e. the lineages which left no sampled extinct or extant descendant. We used a fixed extant sampling probability *ρ* = 1, corresponding to full sampling of all extant lineages.

In order to evaluate precisely the accuracy of the MTFBD inference under different conditions, we simulated phylogenies under four types of rate variation. In condition (1), all rates were constant with a birth rate of λ = 0.15, death rate of *µ* = 0.1 and fossilization rate of *ψ* = 0.1. In condition (2), we simulated two types which differed only by one rate, i.e. a process with two different birth rates or death rates or fossilization rates. In this condition, the other two rates were constant between the two types. In condition (3), we simulated two types which differed by two rates, either both the birth and death rate, or both the birth and fossilization rate. These two rates could be either correlated by type (i.e. the first type had both a low birth rate and a low death rate and the second had both high rates) or reverse-correlated by type (i.e. the first type had a low birth rate and a high death rate, and reverse for the second type). Finally, in condition (4), we simulated a process with four types corresponding to independent variations in both birth and death rates or in both birth and fossilization rates. In all multi-type simulations, a ratio of 6 was used between the low rate value and the high rate value for each rate with variations. These four conditions will allow us to evaluate precisely the accuracy of MTFBD inference when faced with variations in each of the parameter rates, as well as establish whether MTFBD can distinguish between the combined effects of simultaneous variations in several rates.

To ensure that the datasets were composed of phylogenies of similar size and make comparisons easier, the simulated phylogenies were discarded if the number of simulated fossil samples was not between 80 and 120 samples. In addition, multi-type phylogenies were discarded if either type was represented by less than 10% of the final samples (extant and extinct combined), or if the total number of type changes throughout the tree was above 15. The overall type change rate was calibrated depending on the dataset to minimize simulation rejection. A full list of the parameters used for the simulated datasets can be found in Table S2. Simulations were repeated until we obtained 50 replicates for each simulation condition.

#### 2.4.2 Simulating alignments

Molecular sequence alignments of 1000 nucleotides were simulated for all extant samples under an HKY+Γ model with 5 rate categories, discretized from a Gamma distribution with shape parameter *α* = 0.25. Branch rates were simulated under a relaxed uncorrelated clock model. For each tree replicate, the average substitution rate was sampled from a gamma distribution with an expected value of 0.01 substitution per My. Branch-specific rates were then independently drawn under a lognormal distribution with the sampled mean and standard deviation *σ* = 0.1. Molecular sequences were only simulated for the extant samples, and set to unknown (“?”) for the fossil samples.

Morphological character matrices of 300 characters were also simulated under an Mk Lewis model (Lewis, 2001) with two states, with a strict clock and a constant clock rate of 0.1 substitutions per My. Morphological characters were simulated for both extant and extinct samples.

#### 2.4.3 Inference

We ran a full Bayesian phylogenetic inference using the software BEAST2 version 2.6, with the packages MM (morphological models), SA (homogeneous FBD process) and our new version of the MSBD package (multi-type FBD process).

Inferences with fossil samples were run using either the homogeneous FBD process or our new MTFBD model, and using either a total-evidence or a topological constraint approach. The total-evidence approach included both the molecular and the morphological alignment as data, with no additional constraint on ages of the phylogeny. The topological constraint approach used only the molecular sequence alignment, but also constrained fossil samples so they were placed within the correct clade in the extant tree.

In order to evaluate the benefits of including fossils, we ran comparison inferences with no fossil samples under an MTBD model. These inferences were calibrated in time by two different methods, either (a) setting a prior distribution on the MRCA age of the tree, or (b) setting prior distributions on the ages of 5 internal nodes with at least 4 tips, chosen uniformly at random. The prior distribution used for all calibrations was a LogNormal distribution with a standard deviation of 1.0, calibrated so its median corresponded to the true node age and its minimum value was 0.9 times the true node age.

In all inference conditions, the substitution and clock models were set to the models used for simulation for the molecular and (if applicable) morphological data. A wide prior of LogNormal(1.5,2.0) was set on the birth, death and fossilization rates for both the homogeneous and heterogeneous inferences. The priors on the molecular and (if applicable) morphological clock rates were set to Exponential(1.0). All other priors were left to their default distributions.

Inferences were run for a minimum of 100, 000, 000 iterations, or until convergence was reached, with a burn-in period set to 10% of the final chain length. We determined that convergence was reached if an effective sample size (ESS) of minimum 200 was reached on the posterior and clock rate, as calculated by the R package `coda` (Plummer et al., 2006), as well as on the tree topology, as calculated by the R package `RWTY` (Warren et al., 2017).

#### 2.4.4 Output metrics

Since the phylogeny is co-estimated along with the diversification parameters, the birth, death and fossilization rates cannot be meaningfully measured for each edge or node. We thus calculated the relative error and coverage for the estimated birth, death and fossilization rates at each tip (both extant and extinct) of the phylogeny, as well as the average over the entire phylogeny, weighted by the estimated edge lengths. We chose to study the birth and death rates separately rather than e.g. the net diversification rate, in order to evaluate precisely whether the inclusion of fossil samples allows for better estimates of the death rate compared to extant phylogenies. We also measured the relative error and coverage of the divergence times, averaged over all pairs of tips. To measure the error on estimated tree topology, we calculated the average Robinson-Foulds (RF) distance (Robinson and Foulds, 1981) between the phylogenies in the posterior distribution and the true simulated tree. The RF distance was normalized to a scale from 0 to 1. We also built the Maximum Clade Credibility (MCC) summary tree using TreeAnnotator (Bouckaert et al., 2019) and again measured its normalized RF distance to the true phylogeny. The mean RF distance and MCC RF distance were very similar, so we only present this second metric. Finally, in order to compare the MTFBD results to analyses without fossil samples, we re-calculated all measures for the MTFBD analyses on phylogenies including only the extant tips.

For the constant-rate analyses, we measured the relative error using the median of the posterior density, and the coverage using the 95% Highest Posterior Density (HPD) interval as calculated by the R package coda. For the multi-type analyses, these summary metrics can give unreliable results when the posterior distribution is strongly multimodal (Barido-Sottani et al., 2020b). As a result, we measured the relative error using the mode of the posterior density, as calculated by the R package modeest, and the coverage using the 95% Highest Density Region (HDR) as calculated by the R package hdrcde (Hyndman et al., 2021).

### 2.5 Empirical dataset

We used an empirical dataset of cetaceans, previously analyzed under the OBD process in Andréoletti et al. (2022). The dataset is composed of a molecular sequence alignment and a morphological character matrix. The original molecular data was obtained from Steeman et al. (2009), and comprises 6 mitochondrial and 9 nuclear genes, for 87 of the 89 accepted extant cetacean species. Morphological data was obtained from Churchill et al. (2018), the most recent version of a widely-used dataset first produced by Geisler and Sanders (2003) and contains 327 variable morphological characters for 27 extant and 90 fossil taxa (mostly identified at the species level but 21 remain undescribed). Unlike Andréoletti et al. (2022), our inference cannot use occurrence information, thus we chose to keep our dataset at the species level (rather than the generic level used in the OBDP inference) to use all of the available information in the alignments.

Total-evidence inferences were performed on the combined molecular and morphological dataset under the constant-rate FBD model and under our combined MTFBD model in BEAST2. Following the previous analysis, we partitioned the molecular sequence alignment between mitochondrial and nuclear genes. Each partition was set with a separate *GTR* + Γ substitution model with 4 rate categories and an uncorrelated exponential relaxed clock model. The morphological alignment was partitioned by the number of observed states per character and used an Mk Lewis substitution model, correcting for ascertainment bias. All morphological partitions shared the same strict clock model, again following the previous analysis. The fossil ages were sampled during the inference, using uniform distributions based on the lower and upper age bounds of the fossils stratigraphic age uncertainty from the Paleobiology Database (PBDB) provided in Andréoletti et al. (2022).

We also ran the inference using only the extant samples and the molecular sequence alignment under a regular MTBD model. The substitution and clock models were identical to the models used for the inferences with fossils. Since our simulation results showed few differences between the MTBD inference with multiple calibrations and the MTBD inference with only root calibrations, we ran the inference with a root calibration only. We used two different calibrations on the root age of the tree. The first “wide” calibration was set to allow root ages from 35My to 56My, following the root calibration used in Steeman et al. (2009). The second “narrow” calibration was based on the 95% HPD interval obtained from the FBD and MTFBD inferences, and allowed root ages from 44My to 49.5My. Both calibration methods led to very similar estimates regarding both the tree and the rates, so we only show here the results obtained using the narrow calibration.

Inferences were run for a minimum of 300, 000, 000 iterations, or until convergence was reached, with a burn-in period set to 10% of the final chain length. Similar to the simulation runs, we determined that convergence was reached if an effective sample size (ESS) of minimum 200 was reached on the posterior and clock rate, as calculated by the R package coda (Plummer et al., 2006), as well as on the tree topology, as calculated by the R package RWTY (Warren et al., 2017). Maximum clade credibility (MCC) summary trees were obtained using TreeAnnotator, available as part of BEAST2.

## 3 Results

### 3.1 Simulation study

#### 3.1.1 Comparison with the MTBD model without fossils

Here, we compare the results of the inferences under the MTFBD model and under the MTBD model without fossil samples. Since the MTBD inference only includes information for the extant tips, all measures in this section are calculated based only on the extant phylogeny and samples.

Figure 1 shows the relative error and coverage of the birth and death rates, averaged over the reconstructed extant phylogeny. Figure 2 shows the same relative error excluding the inference using the MTFBD with topological constraints, in order to make the comparison easier. The total-evidence MTFBD inferences perform similarly or better than the inferences without fossils when estimating the birth rates. When estimating the death rates, total-evidence MTFBD performs much better than the inferences without fossils, with the exception of two datasets with variations in both birth and death rates (reverse-correlated and independent), in which it obtains similar accuracy as the MTBD without fossils. However, the MTFBD inference with topological constraints shows high levels of error and low coverage in estimates of both birth and death rates across all datasets except the constant-rate dataset. In general, the MTFBD inference with topological constraints overestimates both birth and death rates (85-95% overestimate on average), whereas the total-evidence MTFBD inference shows little bias. The MTBD inference without fossils tends to underestimate both rates, especially the death rates (75% underestimate on average for the birth rates, 95% underestimate on average for the death rates).

**Figure 1:**
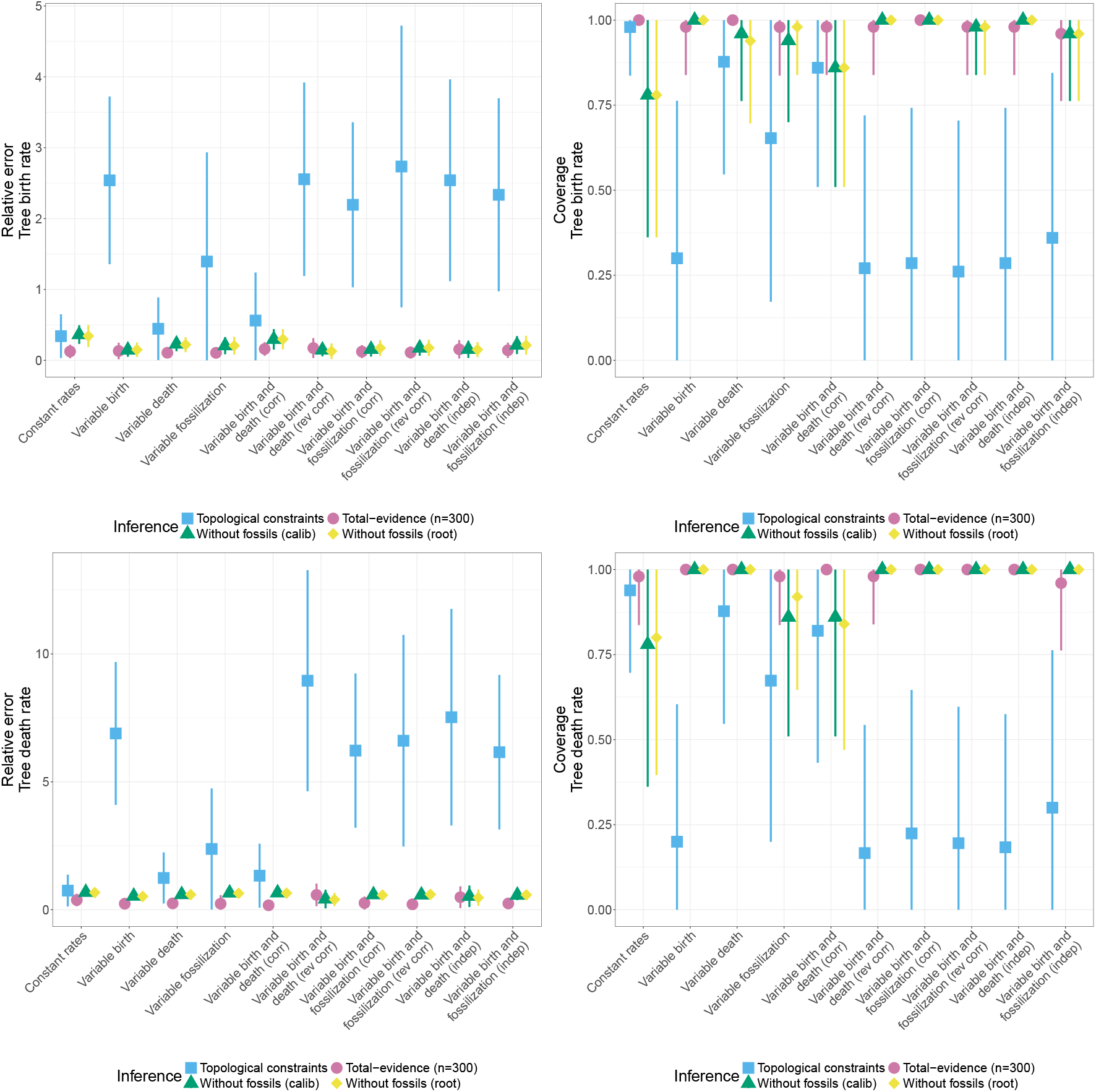
Relative error and 95% HPD coverage of the estimated birth rate (top row) and death rate (bottom row) by inference under the MTBD process without fossil samples (green and yellow) or the MTFBD process (blue and pink), averaged over the phylogeny. The MTFBD inference was run either with topological constraints on the fossil placement (blue) or using a morphological character matrix (pink). The MTBD inference was run with 5 node calibrations (green) or with a calibration on the root age (yellow). The plots show the average and standard deviation over the 50 replicates for each dataset.

**Figure 2:**
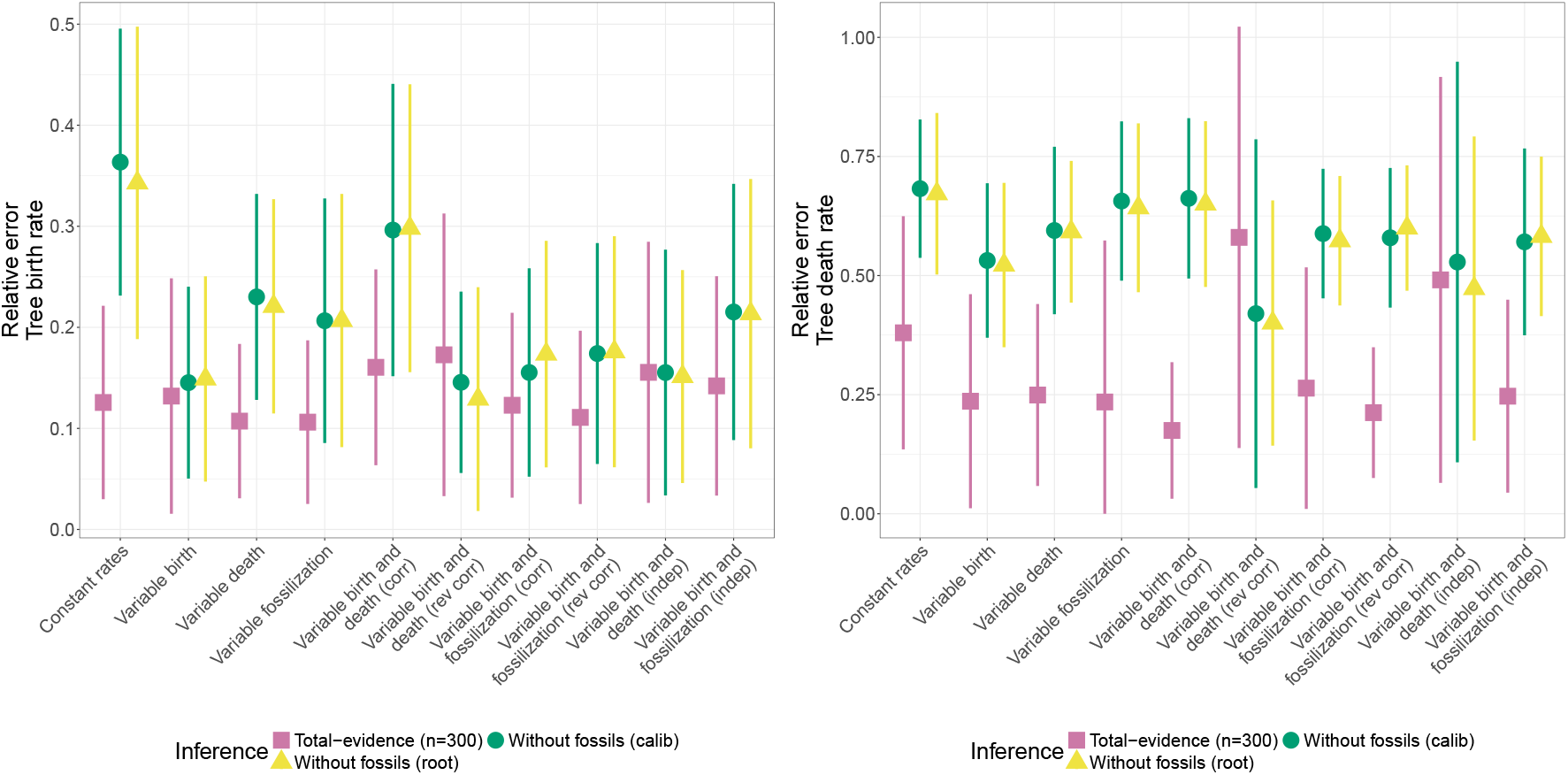
Relative error of the estimated birth rate (left) and death rate (right) by inference under the MTBD process without fossil samples (green and yellow) or the MTFBD process (pink), averaged over the phylogeny. The MTFBD inference was run using a morphological character matrix. The MTBD inference was run with 5 node calibrations (green) or with a calibration on the root age (yellow). The plots show the average and standard deviation over the 50 replicates for each dataset.

Results for the birth and death rates averaged across all tips of the phylogeny are shown in Figures S2 and S3. While these results are broadly consistent with the results averaged across the entire phylogeny, they suggest that individual tip estimates are less reliable overall than estimates over the tree, for both MTBD and MTFBD inferences. However, the metrics calculated on the tips are likely influenced by the presence of type changes very close to the tips, which would change the tip rate recorded by the simulation but would be undetectable by the inference (Barido-Sottani et al., 2020b).

Figure 3 shows the results on the tree topology and divergence time estimates, limited to the extant tree. Divergence times estimates are well estimated by all inferences, however the relative error is lower under the total-evidence MTFBD than under the MTBD model without fossil samples, especially in datasets with rate variations. Similar to the rate estimates, the inference with topological constraints performs worse on both the relative error and the coverage. The extant tree topology is also well estimated across all inferences, although the RF distance is lower in inferences using fossils. The RF distance for the inference with topological constraints is almost zero, which is expected as the topological constraints on the fossil samples also restrict the position of the extant samples to correspond to the true phylogeny.

**Figure 3:**
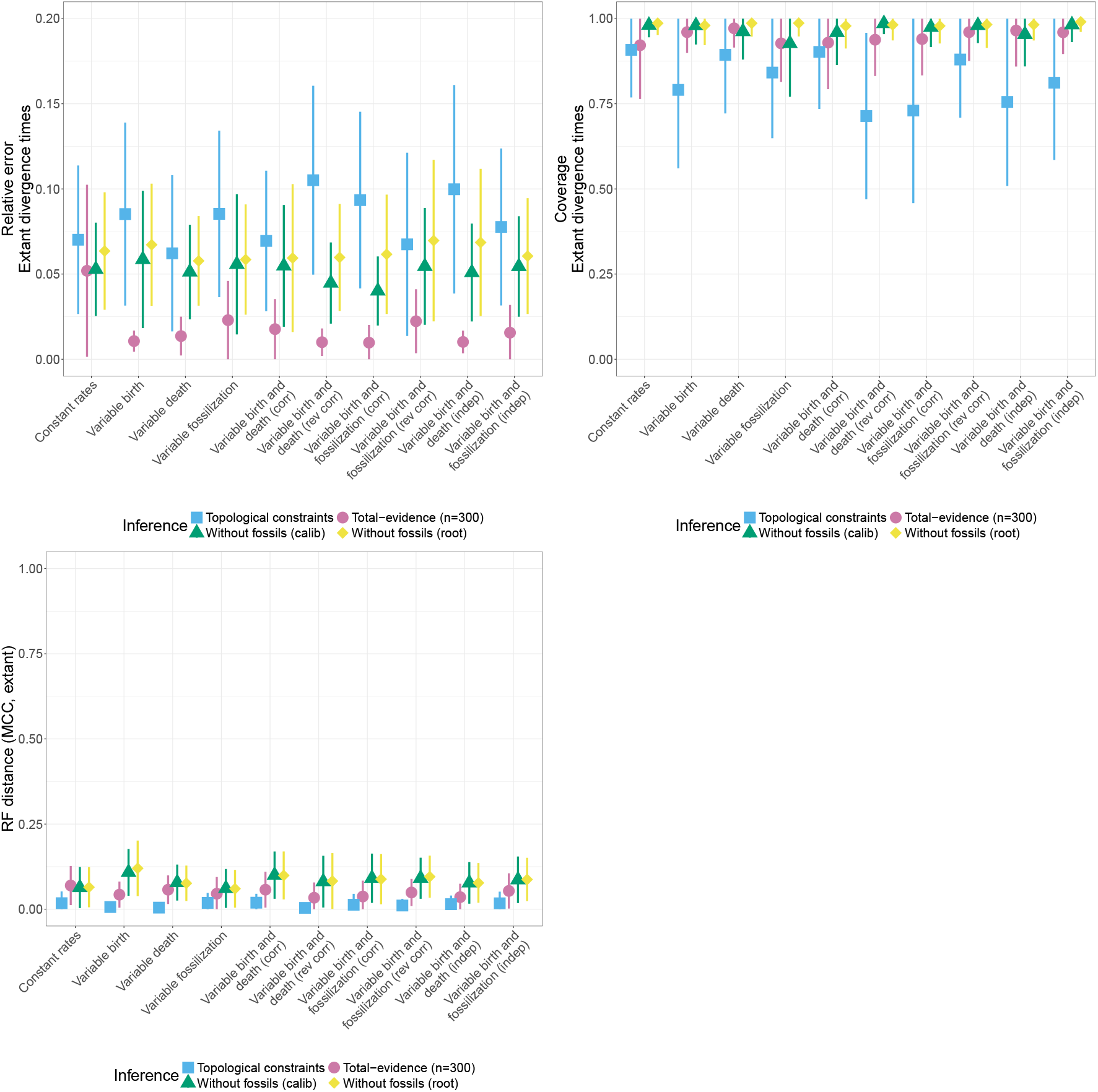
Relative error and 95% HPD coverage of the estimated divergence times, and normalized Robinson-Foulds distance between the extant MCC summary tree and the true extant tree, by inference under the MTBD process without fossil samples (green and yellow) or the MTFBD process (blue and pink). The MTFBD inference was run either with topological constraints on the fossil placement (blue) or using a morphological character matrix (pink). The MTBD inference was run with 5 node calibrations (green) or with a calibration on the root age (yellow). The divergence times measures are averaged over all pairs of tips in the extant phylogeny. The plots show the average and standard deviation over the 50 replicates for each dataset.

Overall, the MTFBD inference with topological constraints performs badly on all tested metrics except for the accuracy of the topology estimates. One possible explanation for this pattern is that the inference with topological constraints contains less information to inform the branch lengths of the phylogeny than the total-evidence inference, which may lead to the higher error observed on the divergence times. Since the branch lengths are essential information to accurately estimate diversification rates, this could explain the higher error on the rate estimates. In addition, the MTFBD inference with topological constraints cannot correctly detect whether fossil samples are sampled ancestors (Figure S8), and SAs are known to play a major role in identifying the parameters of the FBD process (Beaulieu and O’Meara, 2023). Exploring these results further (see Figures S6 and S7) shows that the rate and SA estimates from the MTFBD inference with topological constraints are strongly driven by the priors on birth, death and fossilization rates, and that using narrow priors focused on the true values leads to similar accuracy as for the total-evidence MTFBD inference. However, these narrow priors require a large amount of information on the true underlying process of diversification, which is typically not known for empirical datasets.

#### 3.1.2 Comparison with the FBD model with constant rates

Here, we compare the results of the inferences under the MTFBD model and under the FBD model with constant rates. Note that all measures in this section are calculated on the full sampled tree with both extant and extinct tips.

Figure 4 shows the relative error and coverage of the birth and death rates, averaged over the reconstructed phylogeny. Figure 5 shows the same relative error only for the total-evidence inferences, in order to make the comparison easier. Similarly to the results in the previous section, we can see that the MTFBD inference with topological constraints performs quite badly on estimating the rates. It obtains higher relative errors and lower coverage values on both birth and death rates across all datasets. On the other hand, the total-evidence MTFBD inference performs similarly or better and obtains slightly better coverage values than the constant-rate FBD inference at estimating birth and death rates across most datasets. Similar to the results on the extant tree (Figures 1 and 2), the relative error on the death rate estimates is higher for MTFBD than FBD in two datasets with variations in both birth and death rates (reverse-correlated and independent), which suggests that the inference cannot fully identify combined variations in these two parameters. In general, the MTFBD inference with topological constraints overestimates both birth and death rates (85-95% overestimate on average), whereas the total-evidence MTFBD inference shows little bias. The FBD inference tends to underestimate both rates, especially the death rates (75% underestimate on average).

**Figure 4:**
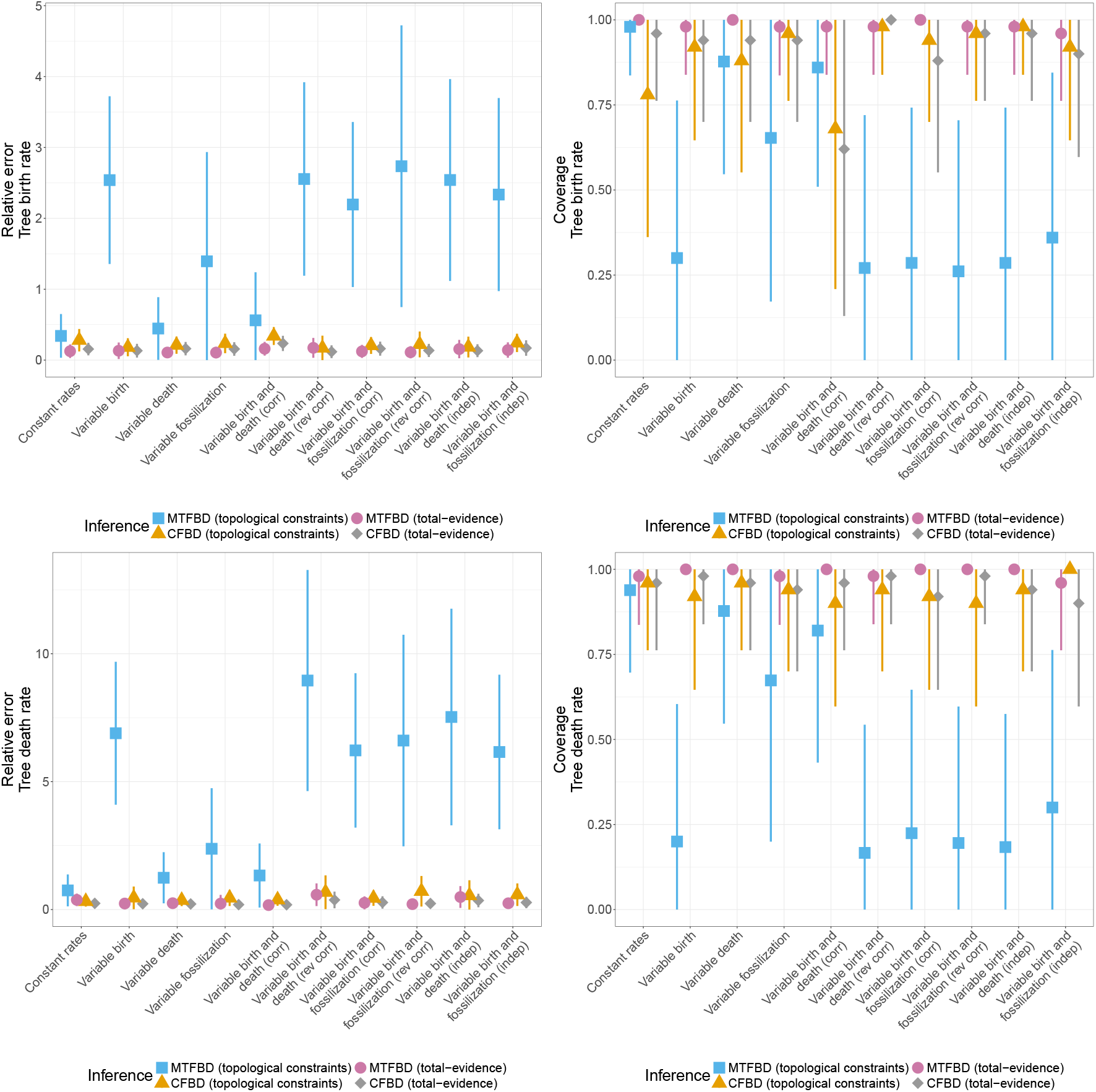
Relative error and 95% HPD coverage of the estimated birth rate (top row) and death rate (bottom row) by inference under a constant-rate FBD process (orange and gray) or the MTFBD process (blue and pink), averaged over the phylogeny. The inference was run either with topological constraints on the fossil placement (orange and blue) or using a morphological character matrix (gray and pink). The plots show the average and standard deviation over the 50 replicates for each dataset.

**Figure 5:**
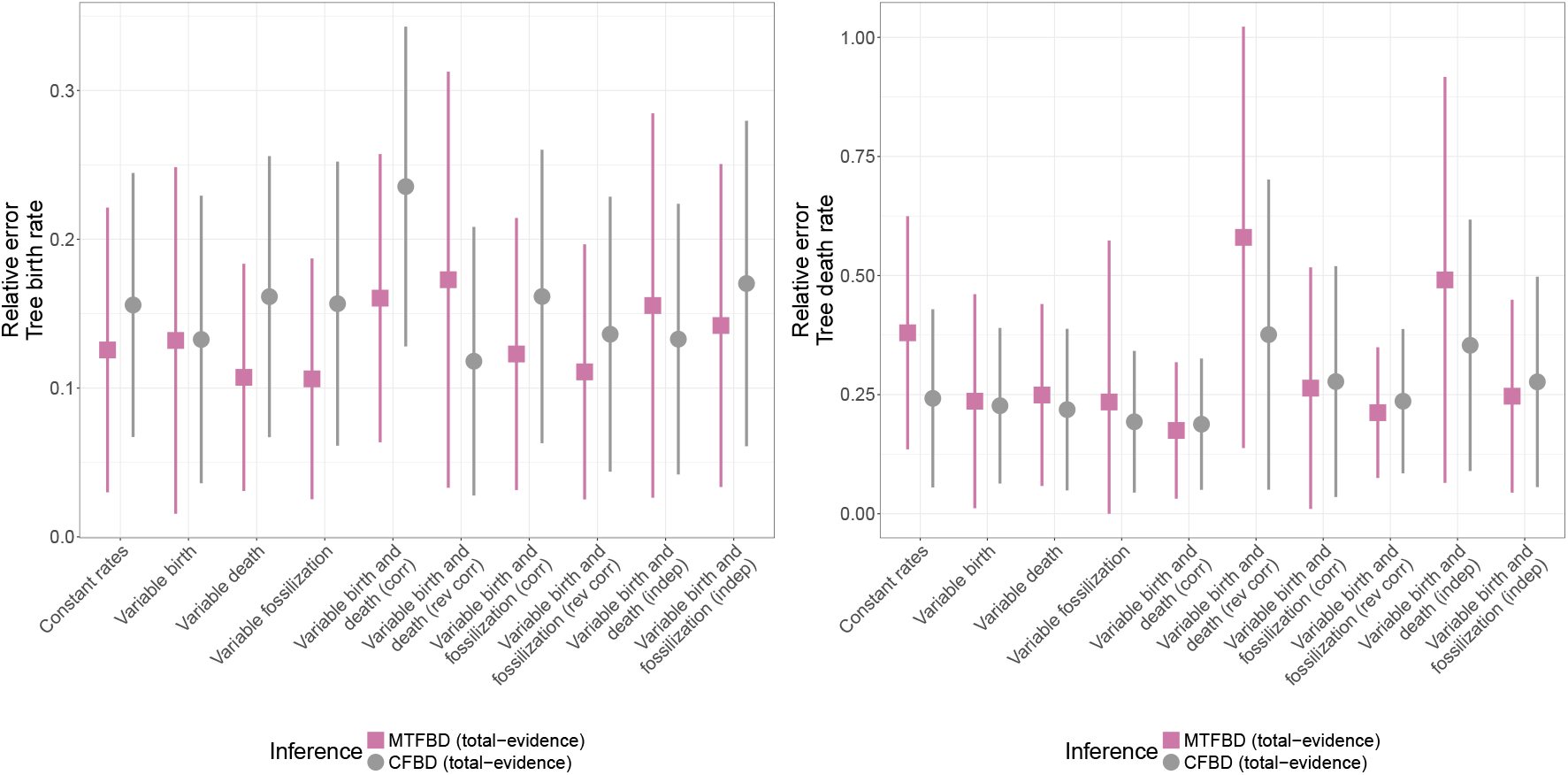
Relative error of the estimated birth rate (left) and death rate (right), by total evidence inference under a constant-rate FBD process (gray) or the MTFBD process (pink), averaged over the phylogeny. The plots show the average and standard deviation over the 50 replicates for each dataset.

Figure 6 shows the relative error and coverage of the fossilization rate, averaged over the reconstructed phylogeny. In this figure, we removed one outlier replicate in the dataset simulated with variable fossilization rate, as its relative error on the fossilization rate was more than 10x the average. We determined that this was due to an artefact of the simulation, which led to two fossil samples with very close ages (difference *<* 0.01) on the same lineages. The fossilization rate was thus estimated to be locally very high on this branch. This illustrates one of the limitations of more complex models such as the MTFBD process, which is the risk of overfitting to noise in the dataset. Aside from this outlier, we can see that both MTFBD and FBD inferences perform well on estimating the fossilization rates, although as before the MTFBD inference with topological constraints presents both higher relative error and lower coverage. The MTFBD inference with topological constraints generally underestimates the fossilization rate (95% underestimate on average), whereas the other inferences show little bias. Finally, we also see higher error and low coverage for the total-evidence MTFBD inference on the dataset in which birth and fossilization rate variations are reversely correlated. One possible explanation is that fossilization rate variations are misidentified in this scenario and attributed to variations in the other rates.

**Figure 6:**
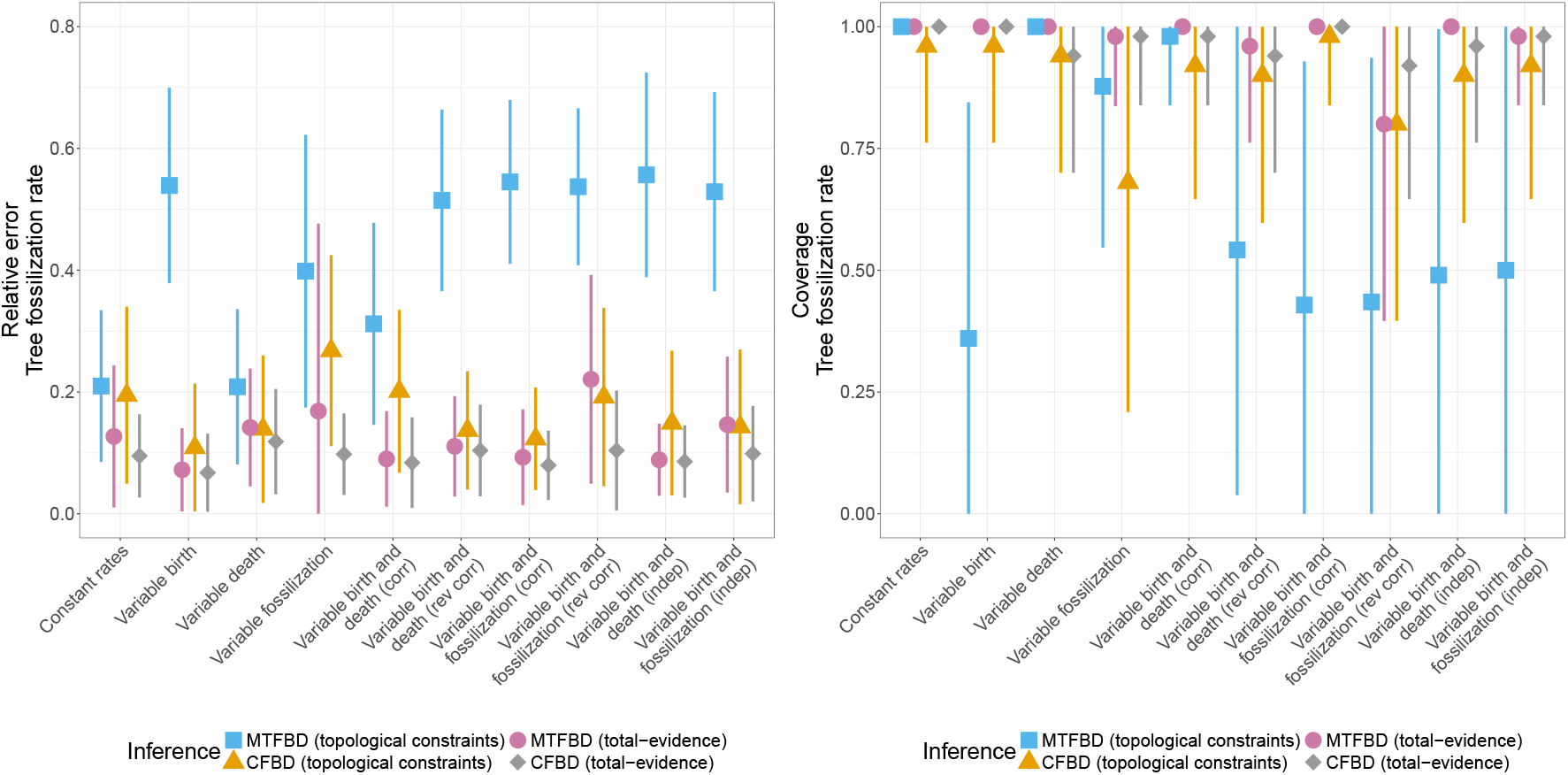
Relative error and 95% HPD coverage of the estimated fossilization rate, by inference under a constant-rate FBD process (orange and gray) or the MTFBD process (blue and pink), averaged over the phylogeny. The inference was run either with topological constraints on the fossil placement (orange and blue) or using a morphological character matrix (gray and pink). The plots show the average and standard deviation over the 50 replicates for each dataset.

Results for the birth, death and fossilization rates averaged across all tips of the phylogeny are shown in Figures S9, S10 and S11. As in the previous section, these results are broadly consistent with the results averaged across the entire phylogeny, but they suggest that individual tip estimates are less reliable overall than estimates over the tree, for both constant-rate FBD and MTFBD inferences. In addition, when a dataset includes variation on a specific rate, FBD inferences show low coverage values for tip estimates for that rate, although the error values remain low. This shows that the FBD inference likely underestimates the uncertainty in estimates, and outputs overly confident credible intervals, possibly due to the simplicity of the model. As before, it is to be noted that the metrics calculated on the tips are likely influenced by the presence of type changes very close to the tips, which would change the tip rate recorded by the simulation but would be undetectable by the inference (Barido-Sottani et al., 2020b).

Figure 7 shows the results on the tree topology and divergence time estimates. We can see that the accuracy is very similar between the MTFBD and the constant-rates FBD inferences, but depends on the method used for placing the fossils, with total-evidence inferences performing better than topological constraints across all datasets. This is coherent with the fact that total-evidence analyses contain more information on the fossils, especially regarding the branch lengths, whereas the analyses with topological constraints only contain information about selected clades. In particular, our topological constraints only assigned fossil samples to their closest extant relative, and thus left a lot of uncertainty on the relative position of fossil samples within each subclade. One thing to note is that the inferences presented here show little uncertainty on the phylogeny estimates, especially for the total-evidence analyses, which could explain why the influence of the tree prior (CFBD or MTFBD) is low. It is possible that the choice of diversification model would make a larger difference if there was less information present in the data, for instance with a much smaller morphological character matrix.

**Figure 7:**
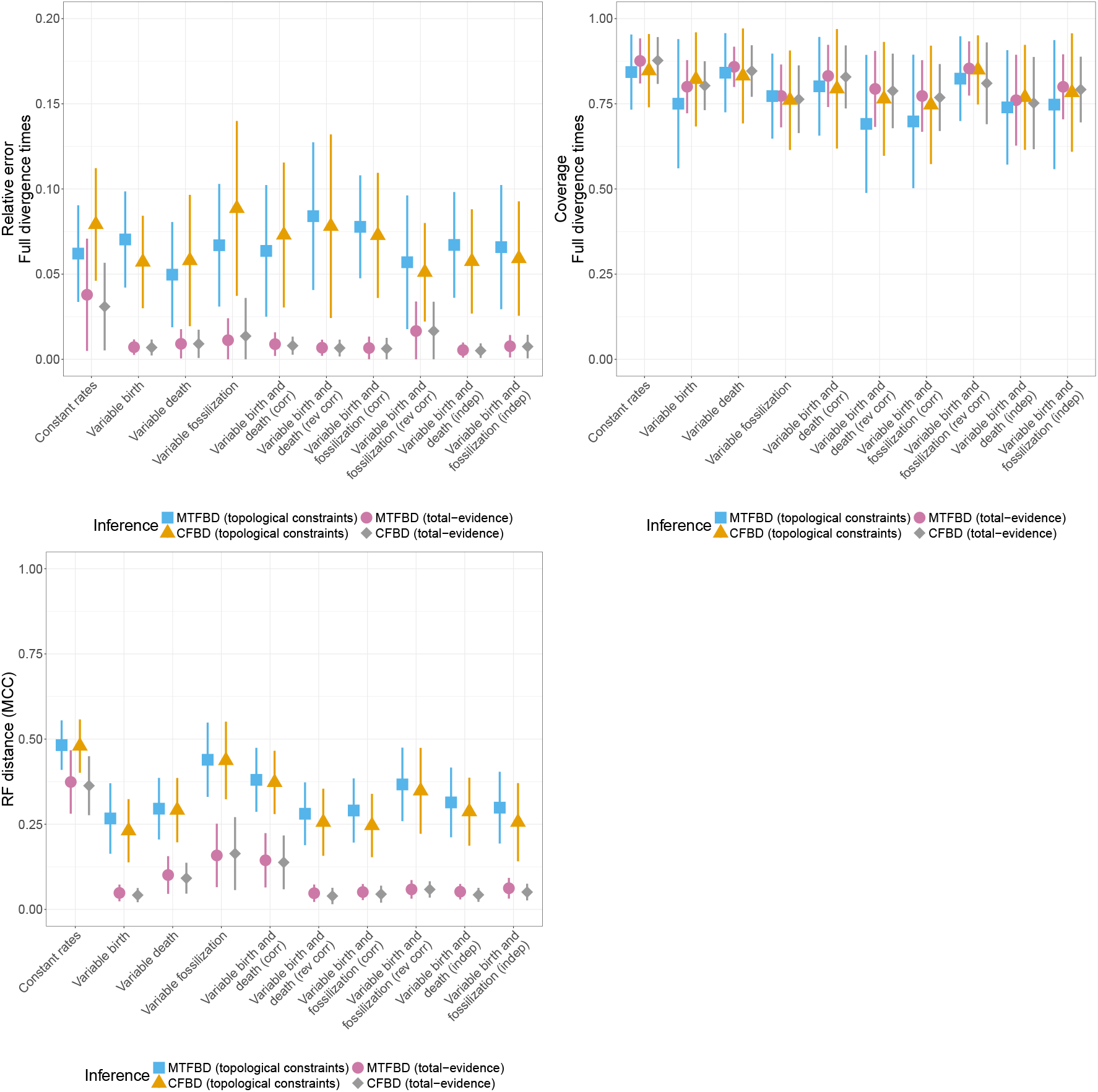
Relative error and 95% HPD coverage of the estimated divergence times and normalized Robinson-Foulds distance between the MCC summary tree and the true tree, by inference under a constant-rate FBD process (orange and gray) or the MTFBD process (blue and pink). The inference was run either with topological constraints on the fossil placement (orange and blue) or using a morphological character matrix (gray and pink). The divergence times measures are averaged over all pairs of tips in the phylogeny. The plots show the average and standard deviation over the 50 replicates for each dataset.

### 3.2 Empirical dataset

Figures 8 and 9 show side-by-side comparisons of the MCC summary tree estimated using either the totalevidence MTFBD model, or the MTBD model without fossil samples, coloured by the mode of the estimated birth or death rate per edge. Figure S12 shows the summary tree obtained under the constant-rate FBD process. Finally, Figure S13 shows the MCC summary tree estimated using the MTFBD model coloured by the mode of the estimated fossil sampling rate per edge.

**Figure 8:**
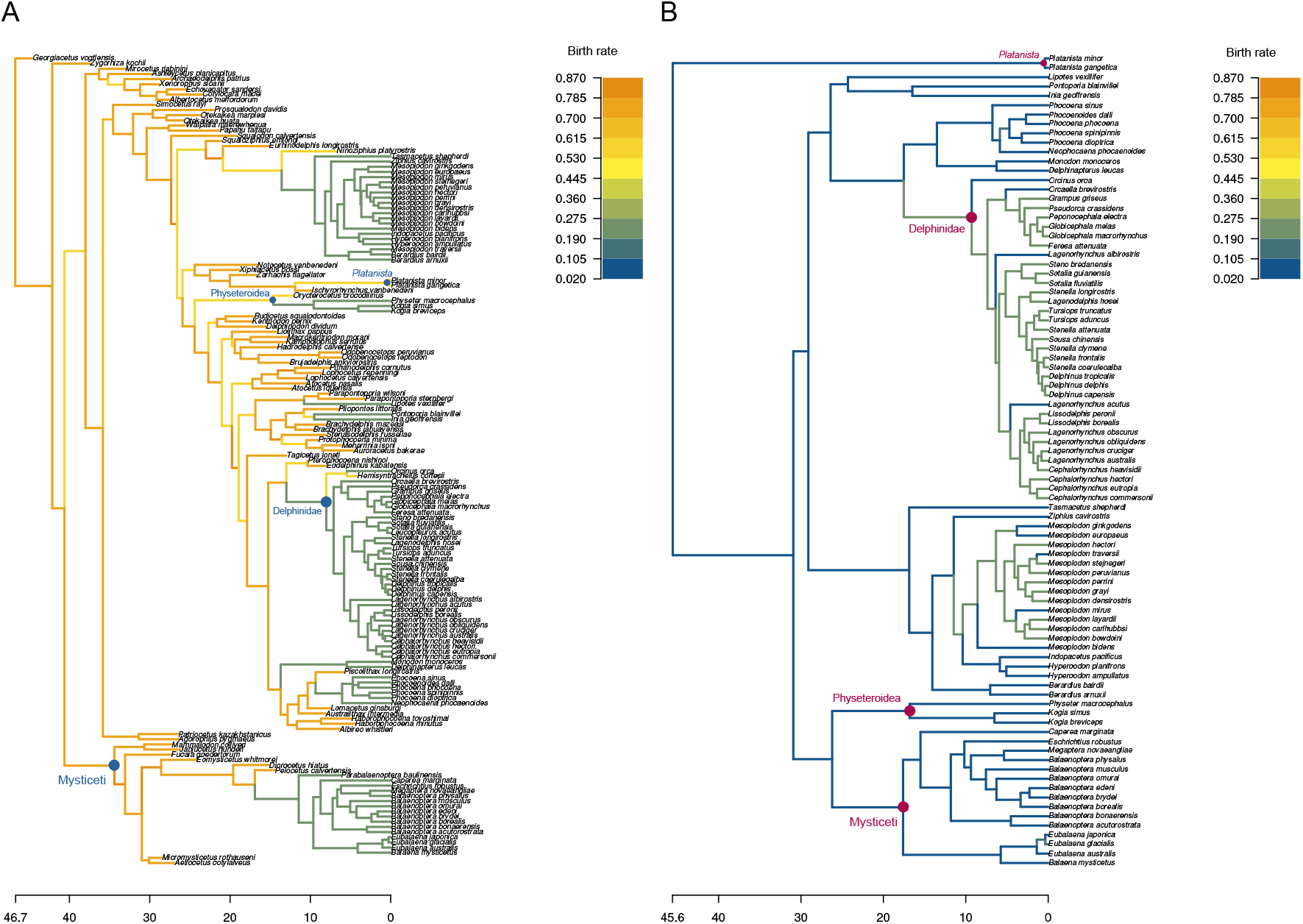
MCC summary tree of the cetaceans phylogeny inferred under a combined MTFBD model (A), or under the MTBD model without fossils (B). Each edge is coloured by the mode of the estimated birth rate for that edge. The MRCA of clades mentioned in the text is indicated by blue (A) or pink (B) dots.

**Figure 9:**
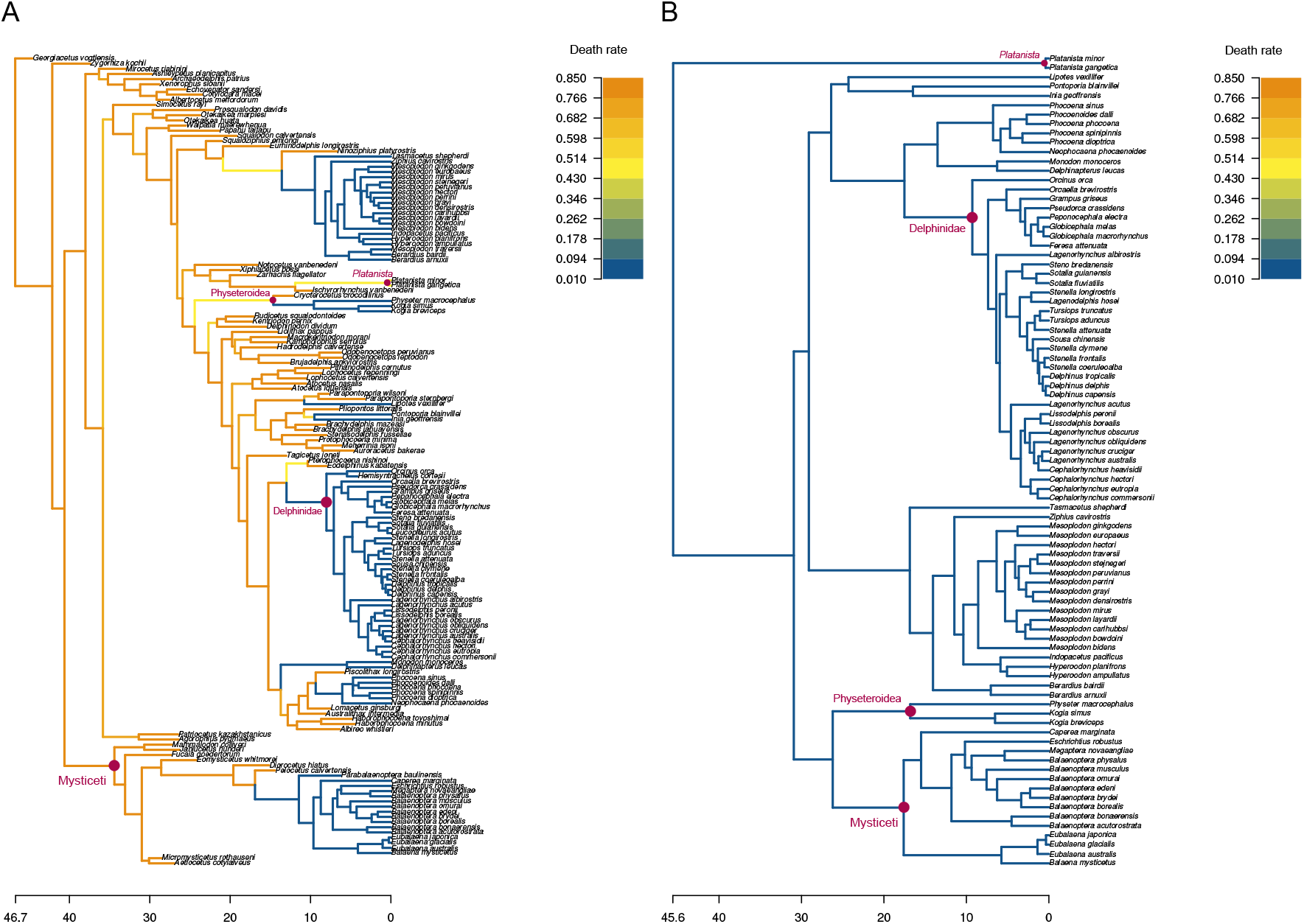
MCC summary tree of the cetaceans phylogeny inferred under a combined MTFBD model (A), or under the MTBD model without fossils (B). Each edge is coloured by the mode of the estimated death rate for that edge. The MRCA of clades mentioned in the text is indicated by pink dots.

First, we compare the MTFBD and FBD inferences. These two inferences estimate a very similar phylogeny (Figures 8 and S10), with an RF distance of 0.175 between the two MCC trees. The mean relative difference of the divergence times between the MTFBD and FBD summary trees is calculated as 0.026, confirming the similarity of the two phylogenies. The median root age is also similar, inferred to 46.8 Ma [44.1 - 49.4] under the MTFBD process and 47.1 Ma [44.5 - 49.9] under the FBD process. The median estimated rates for the constant-rate process are 0.56 for the birth rate, 0.54 for the death rate and 0.022 for the fossil sampling rate. These rates are all coherent with the range of rates estimated under the MTFBD process, however the MTFBD process also identifies large variations between the rates for different clades. In particular, birth, death and fossilization rates are all estimated to much larger values in the backbone of the tree compared to the subclades containing most of the extant species (see Figure S13 for fossilization rate estimates). Within the extant clades, the *Platanista* genus is estimated to have a higher birth rate than the other extant families.

Comparing now the MTBD inference without fossils to the MTFBD inference, we find that including the fossils leads to a quite different phylogeny estimate, with an RF distance of 0.238 and a mean relative difference of the divergence times of 0.223 between the MTBD summary tree and the extant MTFBD summary tree. In particular, the placement of the *Platanista* genus and the Physeteroidea family in the MTBD summary tree are in contradiction both with the other inferences and with the accepted taxonomy of the clade. This is consistent with the simulation results, which show that the tree estimate is more reliable when using fossils. The low performance of the MTBD model in this dataset could also be due in part to our choice of using only one calibration node at the root, which may increase the uncertainty in both tree topology and ages. The original analysis of the molecular dataset in Steeman et al. (2009) did not use fossils, but used a total of 7 calibration nodes. Overall, the birth rates estimated by the MTBD process for the extant species are lower than the birth rates estimated when including fossils, and the death rates are slightly lower than the MTFBD estimates. This is coherent with our simulation results, which showed that the inference without fossils tends to underestimate both birth and death rates, while the MTFBD inference shows little bias. The Delphinidae family is estimated to have a higher birth rate than other extant clades in the MTBD inference, but not the MTFBD inference. Finally, the MTBD inference shows very little heterogeneity in the death rates in the extant clades, while the MTFBD inference identifies several extant lineages with elevated death rate (*Platanista* genus).

## 4 Discussion

We implement here a combined multi-type fossilized birth-death process, which can model variations across lineages for birth, death and fossil sampling rates, while integrating both extant and extinct samples. We demonstrate that total-evidence MTFBD inferences combine the advantages of both the original models, by obtaining more accurate tree estimates than inferences without fossil samples, while providing a full accounting of rate variations between different lineages of the phylogeny. In addition, death rates estimated by the MTFBD inference are more accurate than estimates obtained from inferences without fossil samples, confirming that extinct diversity plays a major role in correctly reconstructing diversification and particularly extinction through time. Through the MTFBD model, phylogenetic inferences can thus obtain a full representation of variations in evolutionary processes, including both extant and extinct information. The MTFBD model is implemented in BEAST2 and can be used either on a fixed phylogeny, or to co-estimate the phylogeny and the diversification parameters, thus providing a powerful framework to account for both rate heterogeneity and tree uncertainty. It is fully integrated into the BEAUti graphical interface and can be combined with all other components of the BEAST2 framework, including prior distributions as well as substitution and clock models.

One crucial finding from our simulations is that MTFBD inferences using topological constraints are inaccurate, and in many cases perform worse than MTBD inferences without fossil samples. One likely explanation for this result is that the inference of rates under a birth-death model is primarily driven by the pattern of branch lengths throughout the phylogeny, and thus depends on the accuracy of these estimates. Unlike total-evidence approaches which include a morphological character matrix and clock model, the topological constraint approach provides no information on the branch lengths of edges leading to the fossil samples. Since the MTFBD process allows each edge to evolve under its own set of parameters, there is thus little information in this inference on the rates associated with fossil parts of the phylogeny. Indeed, our results show that both MTFBD and FBD inferences with topological constraints struggle to estimate the full tree topology, and the MTFBD inference is particularly inaccurate when estimating whether fossil samples represent ancestors to other samples or sister species. The results of the MTFBD inference with topological constraints appear to be largely driven by the prior, and using narrow priors focused on the true rate values leads to similar accuracy as for the total-evidence MTFBD inference. However, setting narrow priors when performing an inference from empirical data is not advisable, as this would require having information on the true underlying process of diversification, which is precisely part of the unknown information we are trying to infer. We therefore advise to use total evidence rather than topological constraints in FBD inferences when possible. If topological constraints are used, we advise to check the robustness of the inference to rate priors, as well as the proportion of fossils inferred to be sampled ancestors. A strong sensitivity to the priors suggests to interpret both the tree and the rate estimates with great caution, and even to give more confidence in an extant-only inference than an inference with fossils. We note that our total-evidence approach uses a large character matrix (300 characters), simulated under a clock model matching perfectly the inference model, and so represents a best-case scenario for this type of inference. Overall, the accuracy of MTFBD inferences is likely to be very dependent on the information available to place fossil samples.

Our simulation results are consistent with previous findings by Mitchell et al. (2019), who found that including fossil samples in multi-type birth-death processes led to improvements in the accuracy of rate estimates across the phylogeny. This contradicts with findings from Beaulieu and O’Meara (2023), who saw no improvements in rate estimates when including fossil samples. One possible contributing factor to this discrepancy is that the MiSSE model uses a fixed number of types, unlike our MTFBD model and fossil-BAMM which both estimate this parameter. Another potential explanation is that Beaulieu and O’Meara (2023) focused on the net diversification rate and extinction fraction rather than the birth and death rates used by our study. This alternative parametrization could contribute to “hide” some of the improvement in estimates, which was focused on death rates in both our results and Mitchell et al. (2019). Both these previous studies tested inferences run on a fixed phylogeny, which was set to the true simulated tree, whereas our MTFBD model is designed for joint analyses estimating both the phylogeny and the diversification process from molecular and/or morphological information. In this situation, we demonstrate that the inclusion of fossil samples in a rate-heterogeneous total-evidence analysis also improves the accuracy of the estimated tree and ages. Since the placement of fossil species is uncertain in many empirical datasets, particularly for the stem group, we believe that the presence of phylogenetic uncertainty represents a more realistic scenario. Beaulieu and O’Meara (2023) also shows that undersampling SAs leads to large decreases in the accuracy of birth and death rate estimates. Similarly, our results demonstrate that the correct inference of the number of SA in the phylogeny is strongly correlated with the accuracy in estimating the diversification rates.

Our MTFBD inference on the cetacean dataset shows important differences to the previous analysis under the OBD process (Andréoletti et al., 2022). In particular, the OBDP estimated tree ages which were much older than previous estimates from the literature and than our MTFBD estimate. Andréoletti et al. (2022) explained these much older ages recovered for this dataset by the heterogeneous sampling of occurrences and fossil samples, which could not be modeled under the OBDP. Since the MTFBD model explicitly integrates heterogeneous fossil sampling, and does not use occurrence information, this likely contributes to our age estimates being closer to those from the literature. The MTFBD inference also estimates that all rates decrease towards the present, unlike the OBDP inference, which showed a spike of birth and death rates close to the present time, and a stable fossil sampling rate. Several factors can contribute to this difference. First, we chose to run our MTFBD inference with all sampled species, while the OBDP inference subsampled the dataset to only one representant per genus. Differences in species delimitation between fossil species compared to extant species can lead to taxonomic inflation in the extinct cetacean lineages (Uhen and Pyenson, 2007), and could lead to a large increase in estimated birth and death rates in the past compared to the present rates, as observed in our inference. In addition, many fossil occurrences were sampled during the Neogene period (20 Ma - 3 Ma). Not accounting for these occurrences in the MTFBD inference could lead to a lower estimated diversity and thus rates during this period, which could also contribute to the observed slowdown.

The combined MTFBD model relies on the same assumptions as the constant-rate FBD process on the nature of fossil data, and thus should be expected to suffer from similar limitations. In particular, the FBD process assumes that fossil samples represent individual specimens, rather than being grouped by species. Empirical datasets commonly violate this assumption, and previous work has shown that as a result the birth and death rates estimated by birth-death models on a phylogeny are not directly equivalent to speciation and extinction rates calculated from fossil species directly (Stadler et al., 2018). In violation of the FBD assumptions, our cetacean dataset does not include any fossil specimens of extant species, which likely contributes to the estimated variation in fossilization rates observed between the past and present clades. Total-evidence FBD inferences are also sensitive to the number of morphological characters included in the dataset (Barido-Sottani et al., 2020a). Since the MTFBD inference relies on the accuracy of branch length estimates to correctly infer the different rates, it will likely be even more affected by small morphological matrices than the FBD process, as well as violations of the morphological clock and substitution models.

One of the limitations which appeared in our simulations is the risk of overfitting, which can lead to strong local over- or under-estimates of the parameters due to noise in the dataset. In one of our replicates, the estimate of the fossilization rate showed a very high spike on one edge, which we believe was due to the presence of two almost simultaneous fossil samples in the simulated phylogeny. Overfitting is a common issue with more complex models, especially when available information is limited. One way to reduce this issue is to impose strong priors on the estimated parameters, which will minimize the posterior probability of biologically implausible outcomes. Although this was not a focus of this study, previous work has shown that the results of the MTBD model are sensitive to the prior on the type change rate *γ* (Barido-Sottani et al., 2020b). One of the goals for future work on our model will be to expand the regularization options and priors available to users. For instance, the RevBayes implementation of the multi-type birth-death process uses a joint distribution for the different birth and death rates rather than estimating them all independently (Hö hna et al., 2019). This blocks the inference of separate regimes with very close rates, and in turn limits inference noise. In general, we advise users of Bayesian phylogenetic inference tools to always consider their results critically, especially when using complex models with a large number of parameters.

Another approach to improve the accuracy of inference under the MTFBD model is to increase the amount of information available to the analysis. The empirical dataset we used here was previously analyzed under the OBDP model (Andréoletti et al., 2022), an expanded version of the FBD process which allows for the integration of fossil occurrences, i.e. fossil samples for which only age information is available. Our results show strong differences in the inferred patterns of diversification between the MTFBD estimates and the original OBDP analysis. This comparison is limited by methodological differences between the two analyses, as the OBDP analysis subsampled the dataset to the genus level and did not account for rate heterogeneity across lineages in the phylogeny, whereas the MTFBD analysis omitted the occurrence information. However, the OBDP analysis demonstrates that occurrences provide important information on the total diversity and strongly affect rate estimates. Integrating occurences into the MTFBD model would thus provide a more detailed and accurate representation of the diversification process through time.

Finally, the current inference allows all birth, death and fossilization rates to vary fully independently from each other. However, the cetacean dataset shows strong correlations between rates within lineages, such that lineages show either high or low estimates for all three parameters. Currently it remains unclear whether this pattern reflects a true correlation between the different processes in this clade, or is due to the inability of the model to clearly distinguish between the relative contributions of the different rates. In particular, our simulation results show that the MTFBD inference is less accurate in estimating death rates in datasets with combined variations in both birth and death rates. Similarly, the fossilization rate estimates are slightly less accurate in datasets with combined variations in both birth and fossilization rates. In the future, implementing explicit model selection in the MTFBD model, for instance by allowing only one rate to vary at each regime change point, could help disentangle the different processes and establish how well the true drivers of diversity can be identified.

In conclusion, our MTFBD model opens the way for a more detailed and accurate inference of the heterogeneity of diversification and fossilization processes, fully accounting for both present and past diversity. Future expansions could focus on including more of the available information and more analysis options.

## 5 Acknowledgements

JBS was supported by the European Union’s Horizon 2020 Research and Innovation Programme under the Marie Sklodowska-Curie grant agreement No. 101022928. This work was performed using HPC resources from GENCI–IDRIS (Grants A0130313658 and A0150313658). We thank the Morlon team at IBENS for their very helpful suggestions and feedback on an earlier draft of this manuscript.

## 6 Data availability

The extended model is implemented in the BEAST2 package MSBD v2.0.3 and later. The package can be installed directly through the BEAUti interface or from the Bitbucket repository https://bitbucket.org/bjoelle/msbd/. The R scripts used for simulation, analysis and plotting, the simulated and empirical data files and the XML configuration files used to run BEAST2 can be found in the Zenodo repository https://doi.org/10.5281/zenodo.14794140.

