## Supplementary materials for "Integrating fossils samples with heterogeneous diversification rates: a combined Multi-Type Fossilized Birth-Death model"

### Supplementary Information

#### 1 Validation of the implementation

As described in the main text, we used a combined validation procedure composed of two separate steps, simulation and inference. By assessing the match between the true simulation parameters and the estimates from the inference, this allows us to check that our MTFBD implementation corresponds exactly to the MTFBD model used for simulation. A full list of the distributions used for simulating the validation phylogenies and running the inference can be found in Table 1.

| Parameter | Distribution |
|---|---|
| $t_{origin}$ | Normal($\mu = 1$, $\sigma = 0.2$) |
| $n^*$ | Poisson($\lambda = 4$) |
| $\gamma$ | LogNormal($M = -1.5$, $S = 0.5$) |
| $\lambda$ | LogNormal($M = -1$, $S = 0.5$) |
| $\mu$ | LogNormal($M = -2$, $S = 0.5$) |
| $\psi$ | LogNormal($M = -1$, $S = 0.5$) |

Table 1: List of sampling distributions used in the simulation step of the validation procedure. The same distributions were used as priors in the inference step. The sampling proportion at present was fixed to $\rho = 1$ in all simulations.

The results of the validation procedure on 1000 replicates are shown in Figure 1. We tested as parameters the type change rate $\gamma$, the total number of types $n^*$, the mean birth ($\lambda$), death ($\mu$) and fossilization ($\psi$) rates averaged over all tips, and the origin time $t_{origin}$. For all tested parameters, the proportion of replicates where the true simulation value falls into the $\alpha$% highest posterior density (HPD) credible interval closely follows $\alpha$, showing that the simulation and inference implementations match.

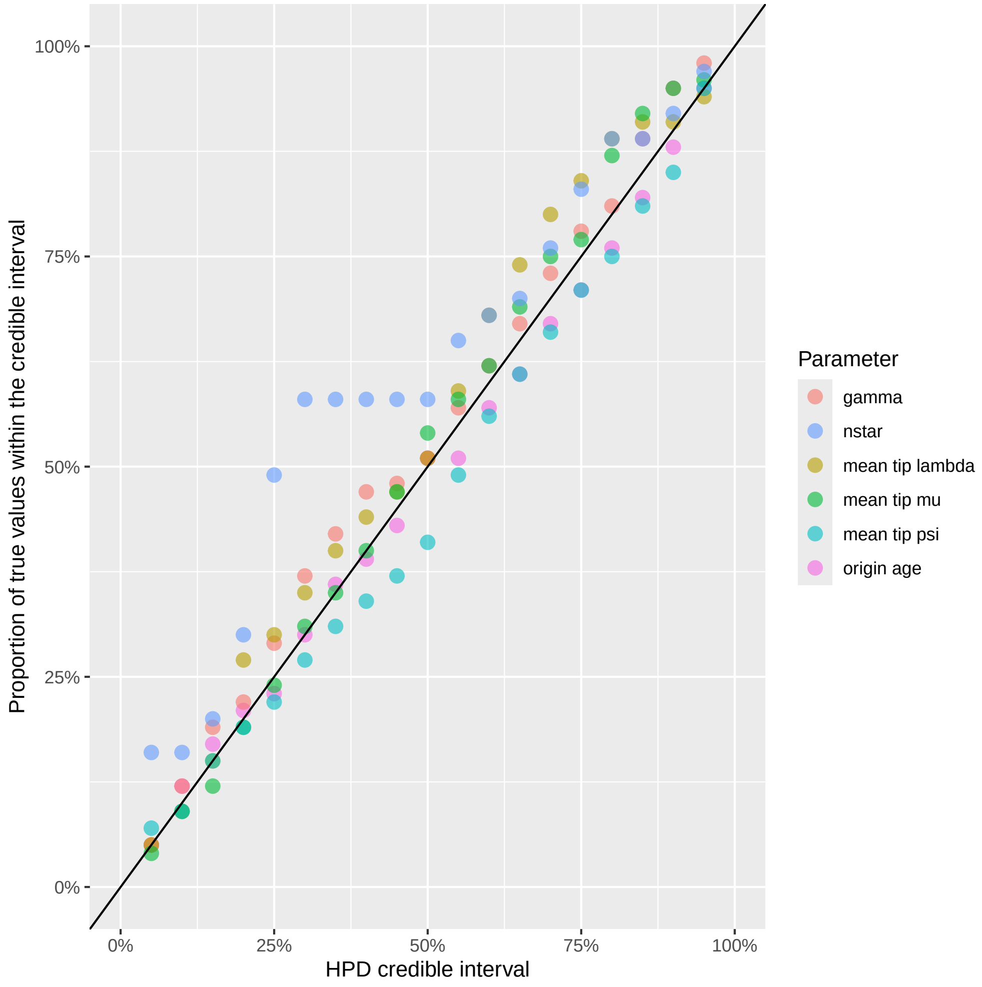


Figure 1: Proportion of the true parameter values used for simulation which fall into the HPD interval for different values of $\alpha$. The parameters shown are (in the order of the legend) the type change rate $\gamma$, the total number of types $n^*$, the mean birth ($\lambda$), death ($\mu$) and fossilization ($\psi$) rates averaged over all tips, and the origin time $t_{origin}$. All proportions are averaged over 1000 simulation replicates. The identity line 1:1 is plotted in black. Note that the total number of types $n^*$ is an integer parameter with discrete values, so the calculated proportions show jumps instead of a continuous pattern.

### 2 Simulation study

#### 2.1 Parameters used for simulation

The parameter used for simulating the phylogenies under the MTFBD process are shown in Table 2. Note that correlated and reverse correlated datasets contains two types, type 1 ($\lambda_1$, $\mu_1$, $\psi_1$) and type 2 ($\lambda_2$, $\mu_2$, $\psi_2$), whereas independent datasets contains four types, which include all possible combinations of rates shown in the table. Parameter values were calibrated to obtain trees of similar sizes and origin times, with at least 10% of samples from each type in the process.

| Dataset | Birth rate(s) | Death rate(s) | Fossilization rate(s) | Type change rate |
|---|---|---|---|---|
| Constant rates | $\lambda = 0.15$ | $\mu = 0.1$ | $\psi = 0.1$ | $\gamma = 0$ |
| Variable birth | $\lambda_1 = 0.05, \lambda_2 = 0.3$ | $\mu = 0.1$ | $\psi = 0.5$ | $\gamma = 0.2$ |
| Variable death | $\lambda = 0.15$ | $\mu_1 = 0.03, \mu_2 = 0.18$ | $\psi = 0.2$ | $\gamma = 0.05$ |
| Variable fossilization | $\lambda = 0.15$ | $\mu = 0.1$ | $\psi_1 = 0.05, \psi_2 = 0.3$ | $\gamma = 0.05$ |
| Variable birth and death (correlated) | $\lambda_1 = 0.05, \lambda_2 = 0.3$ | $\mu_1 = 0.03, \mu_2 = 0.18$ | $\psi = 0.2$ | $\gamma = 0.05$ |
| Variable birth and death (reverse correlated) | $\lambda_1 = 0.05, \lambda_2 = 0.3$ | $\mu_1 = 0.18, \mu_2 = 0.03$ | $\psi = 0.5$ | $\gamma = 0.1$ |
| Variable birth and fossilization (correlated) | $\lambda_1 = 0.05, \lambda_2 = 0.3$ | $\mu = 0.1$ | $\psi_1 = 0.1, \psi_2 = 0.6$ | $\gamma = 0.2$ |
| Variable birth and fossilization (reverse correlated) | $\lambda_1 = 0.05, \lambda_2 = 0.3$ | $\mu = 0.1$ | $\psi_1 = 0.9, \psi_2 = 0.15$ | $\gamma = 0.2$ |
| Variable birth and death (independent) | $\lambda_1 = 0.05, \lambda_2 = 0.3$ | $\mu_1 = 0.03, \mu_2 = 0.18$ | $\psi = 0.5$ | $\gamma = 0.05$ |
| Variable birth and fossilization (independent) | $\lambda_1 = 0.05, \lambda_2 = 0.3$ | $\mu = 0.1$ | $\psi_1 = 0.1, \psi_2 = 0.6$ | $\gamma = 0.2$ |

Table 2: Simulation parameters used for the simulated datasets.

#### 2.2 Comparison with the MTBD model without fossils

Figure 2 shows the relative error and coverage of the birth and death rates, averaged over all extant tips. Figure 3 shows the same relative error excluding the inference using the MTFBD with topological constraints, in order to make the comparison easier. We observe similar patterns as the metrics calculated on the extant phylogeny (see Main Text, Figures 1 and 2), with some exceptions. First, the coverage of the MTFBD inference with topological constraints is much better for the tips on both birth and death rates, and similar to the coverage obtained with the other inferences. We also see higher error and lower coverage for the other types of inferences for both birth and death rates. These results suggest that the inference on specific tips is less reliable than for the global pattern across the entire tree. One possible explanation is that tip rates can be driven by type changes which are very close to the tips, and thus not identifiable from the phylogeny (as was previously observed in Barido-Sottani et al. (2020)).

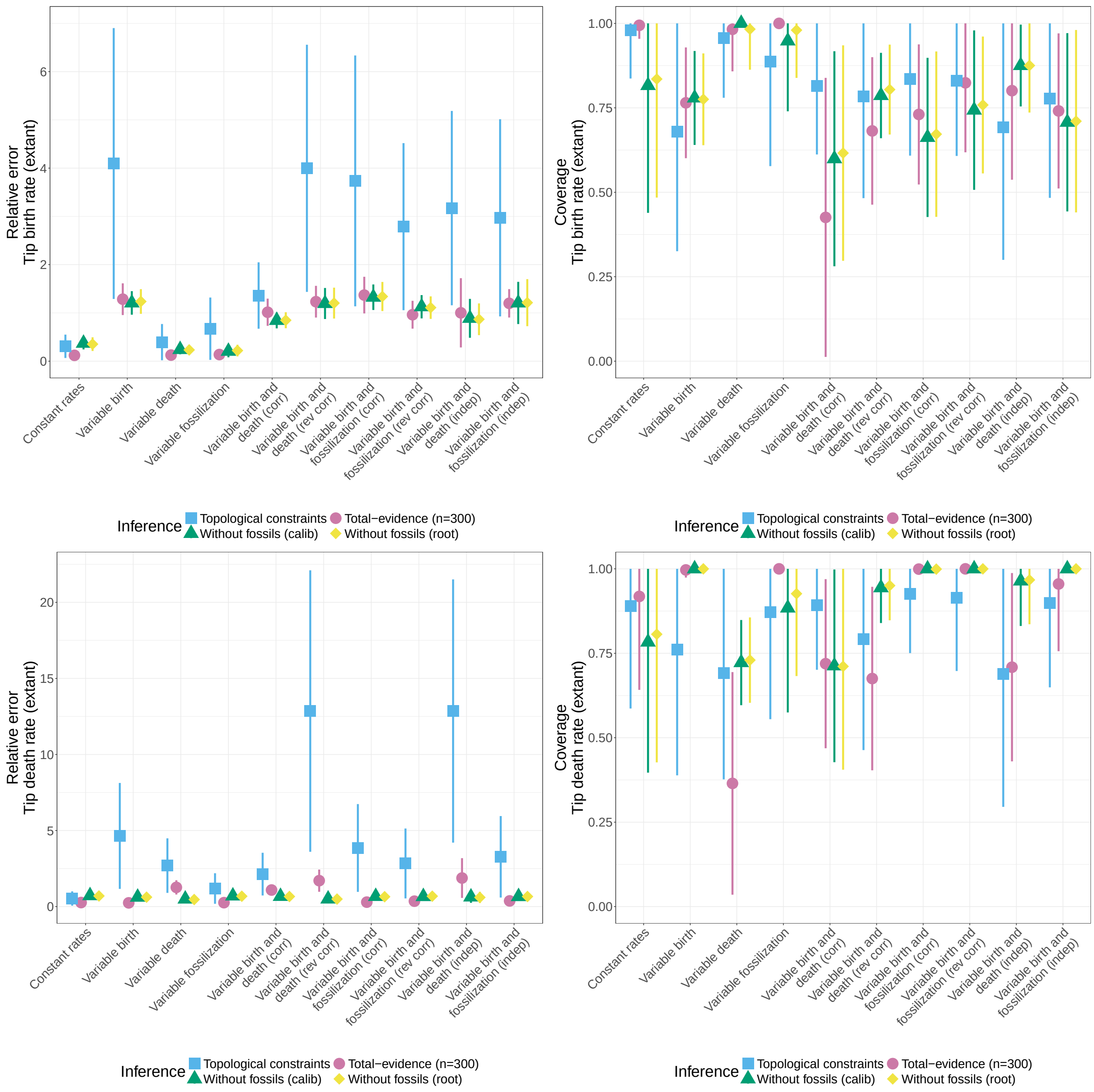


Figure 2: Relative error and 95% HPD coverage of the estimated birth rate (top row) and death rate (bottom row), by inference under the MTBD process without fossil samples (green and yellow) or the MTFBD process (blue and pink), averaged over all extant tips. The MTFBD inference was run either with topological constraints on the fossil placement (blue) or using a morphological character matrix (pink). The MTBD inference was run with 5 node calibrations (green) or with a calibration on the root age (yellow). The plots show the average and standard deviation over the 50 replicates for each dataset.

Figure 4 shows the accuracy of the MSBD and MTFBD inference on the type change rate $\gamma$. The relative error is quite similar across all methods, however the coverage is generally better for analyses without fossils. The MTFBD analyses have particularly low coverage, in keeping with the low accuracy of this method of inference across the board. The type change rate is consistently underestimated by all inferences, which

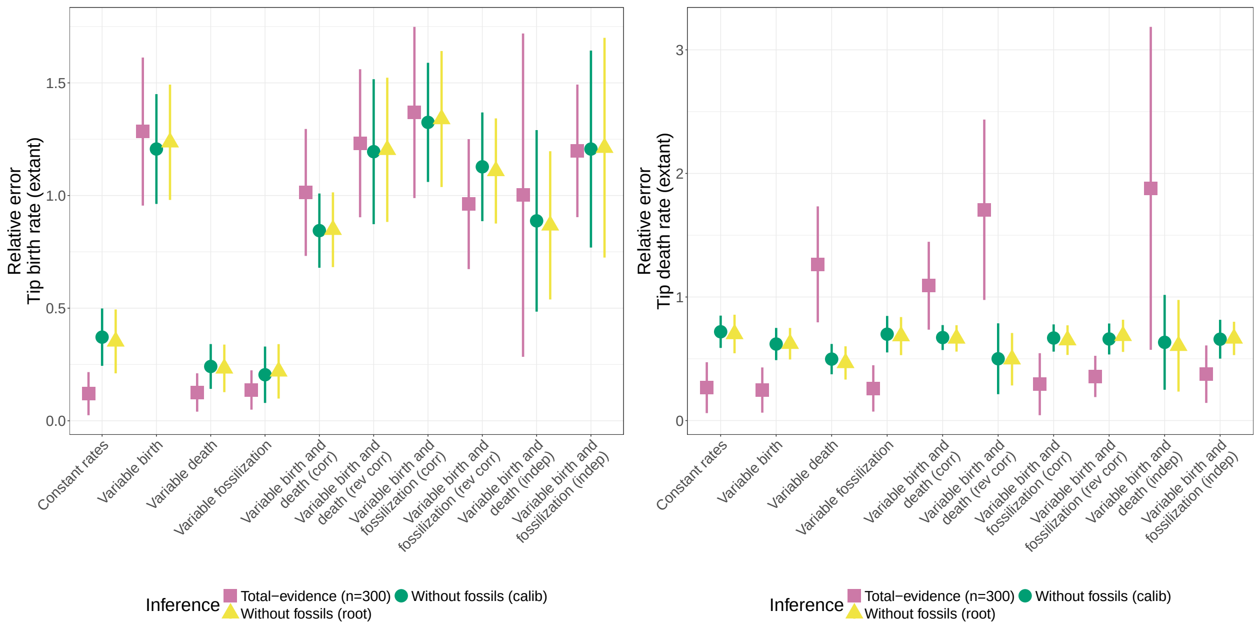


Figure 3: Relative error of the estimated birth rate (left) and death rate (right), by inference under the MTBD process without fossil samples (green and yellow) or the MTFBD process (pink), averaged over all extant tips. The MTFBD inference was run using a morphological character matrix. The MTBD inference was run with 5 node calibrations (green) or with a calibration on the root age (yellow). The plots show the average and standard deviation over the 50 replicates for each dataset.

is logical since the implemented model assumes that there are no type changes in the unsampled parts of the tree, and thus underestimates the number of changes in the process. Overall, the low accuracy of these estimates is consistent with previous results from Barido-Sottani et al. (2020), who found that the type change rate is difficult to estimate accurately and heavily influenced by the prior. Note that the low coverage for the dataset with constant rates is likely due to the true value in that dataset being $\gamma = 0$, a value excluded by the prior on this parameter.

Figure 5 presents the Variation of Information (VI) distance, a measure of the difference between the estimated clustering of tips into types and the true simulated types (Meilă, 2003). This metric shows whether the MSBD and MTFBD inference can accurately identify which samples share the same evolutionary regime, independently of how accurate the parameter estimates for that regime are. The VI distance is also independent of the specific ordering of types. We normalized the VI distance to a range of 0 (identical clusterings) to 1 (fully different clusterings) and averaged it over the posterior distribution. Overall, the VI estimates are quite similar between inferences, although the inference with fossils is more accurate on datasets with variation in the fossilization rate, alone or combined with variation in birth rate. The inference with fossils is also slightly more accurate in the constant-rate scenario.

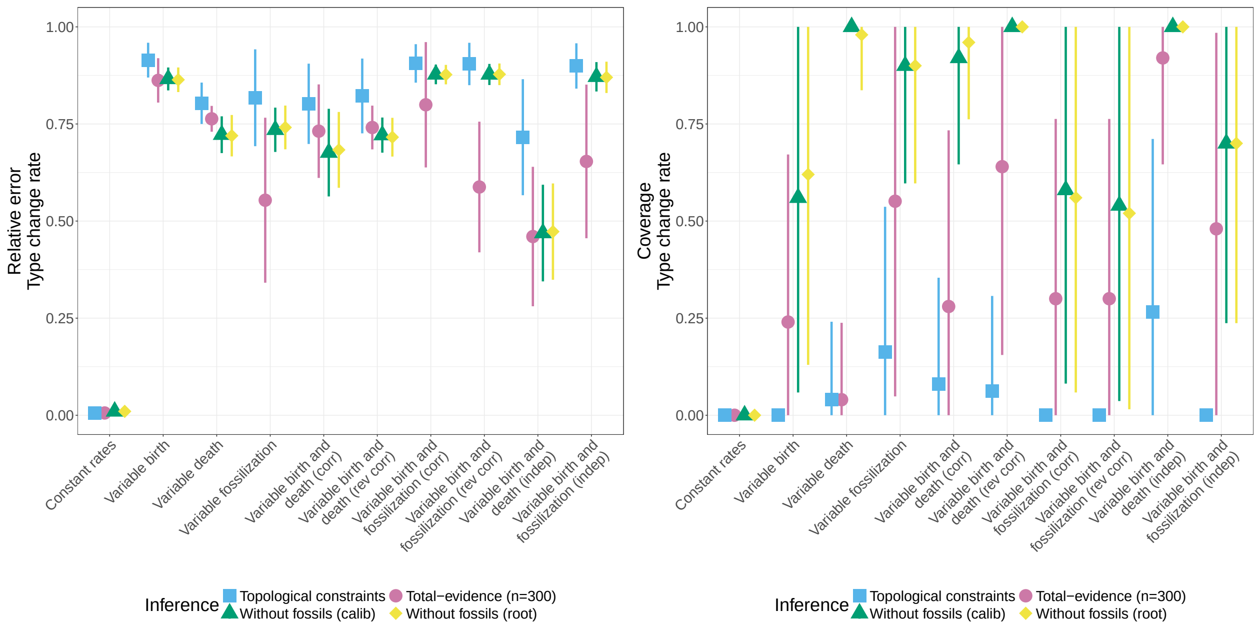


Figure 4: Relative error and 95% HPD coverage of the estimated type change rate, by inference under the MTBD process without fossil samples (green and yellow) or the MTFBD process (blue and pink), averaged over all extant tips. The MTFBD inference was run either with topological constraints on the fossil placement (blue) or using a morphological character matrix (pink). The MTBD inference was run with 5 node calibrations (green) or with a calibration on the root age (yellow). The plots show the average and standard deviation over the 50 replicates for each dataset.

#### 2.3 Further exploration of the MTFBD inference with topological constraints

In order to further analyze the results obtained under the MTFBD model using topological constraints, we checked whether the poor performance of this analysis could be due to the setup of the analysis, namely the lack of SAs in the default starting tree, or the choice of default rate priors. We selected the dataset with variable birth rate and reran the inference either with (1) a starting tree including SAs or (2) narrower priors on the type change rate and the birth, death and fossilization rates, focused around the true parameter values.

Results are shown in Figures 6 and 7 for the rate estimates measured across the phylogeny, as well as for the estimated number of SAs. We can first see that the choice of starting tree does not influence the results. This is expected if our inference has converged, and confirms that the underestimate of the number of SAs by the MTFBD inference with topological constraints is not an artefact of the implementation. Second, we observe that setting narrower priors results in much better estimates of the birth, death and fossilization rates, comparable to the results obtained with the total-evidence MTFBD analysis. The error on the estimated number of SAs also decreases sharply when using the narrower priors on the MTFBD inference with topological constraints. Overall, these results show that estimates under this type of inference

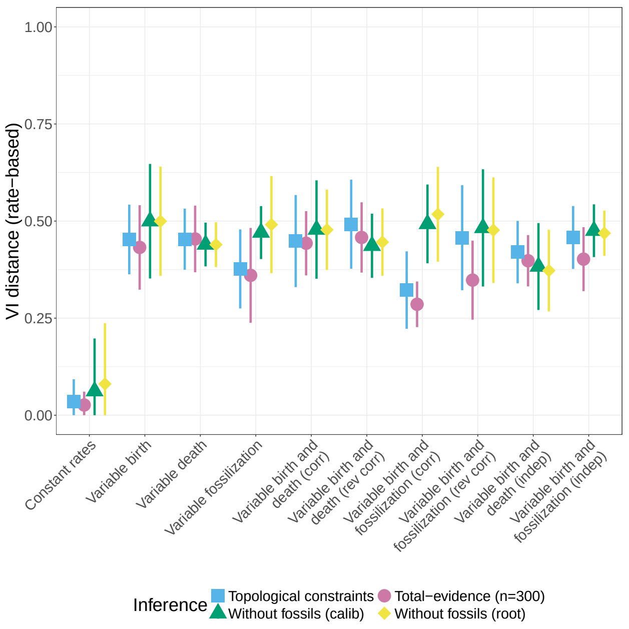


Figure 5: Average normalized VI distance between the estimated clustering of tips into types and the true clustering, by inference under the MTBD process without fossil samples (green and yellow) or the MTFBD process (blue and pink), averaged over all extant tips. The MTFBD inference was run either with topological constraints on the fossil placement (blue) or using a morphological character matrix (pink). The MTBD inference was run with 5 node calibrations (green) or with a calibration on the root age (yellow). The plots show the average and standard deviation over the 50 replicates for each dataset.

are strongly driven by the priors, likely due to the lack of information present in the dataset.

#### 2.4 Comparison with the FBD model with constant rates

Figure 8 shows the relative error and coverage of the number of estimated sampled ancestors (SA) among all fossil tips. We can see that the MTFBD inference with topological constraints performs much worse on both measures. Both total-evidence inferences recover accurately the number of SAs in the phylogeny, although the MTFBD inference shows lower coverage, and more variation between replicates within a dataset. The FBD inference with topological constraints also shows higher error on the number of SAs than both total-evidence inferences, although to a lesser extent than the corresponding MTFBD inference. Overall, these results show a similar pattern as the results obtained on the accuracy of the birth, death and fossilization rates, which is expected as the presence and number of SAs plays a major role in identifying the different rates in the constant-rate FBD process (Beaulieu and O'Meara, 2023).

Figure 9 shows the relative error and coverage of the birth and death rates, averaged over all extant and extinct tips. Figure 10 shows the same relative error only for the total-evidence inferences, in order to make the comparison easier. We observe similar patterns as the metrics calculated on the extant phylogeny (see Main Text, Figures 3 and 4), with some exceptions. First, the coverage of the MTFBD inference with

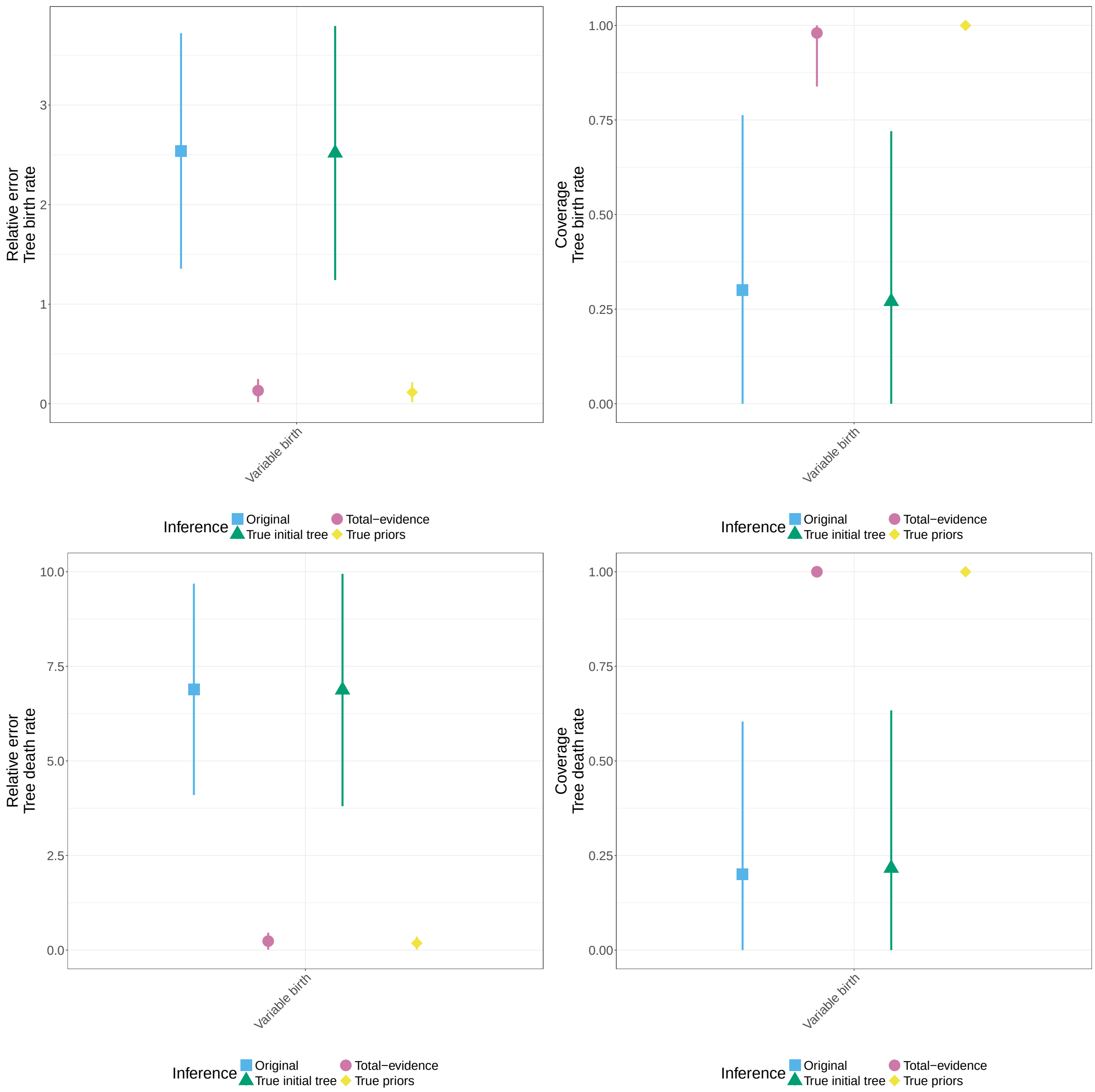


Figure 6: Relative error and 95% HPD coverage of the estimated birth rate (top row) and death rate (bottom row), by inference under the MTFBD process, averaged over the phylogeny. The inference was run either with topological constraints on the fossil placement (blue, green, yellow) or using a morphological character matrix (pink). The plots show the average and standard deviation over the 50 replicates for the dataset with variable birth rate.

topological constraints is much better for the tips on both birth and death rates, and similar to the coverage obtained with the total-evidence MTFBD inference. We also observe low coverage for the FBD inferences on both birth and death rates, particularly for the datasets which contain variation in the corresponding rate.

Figure 11 shows the relative error and coverage of the fossilization rate, averaged over all extant and

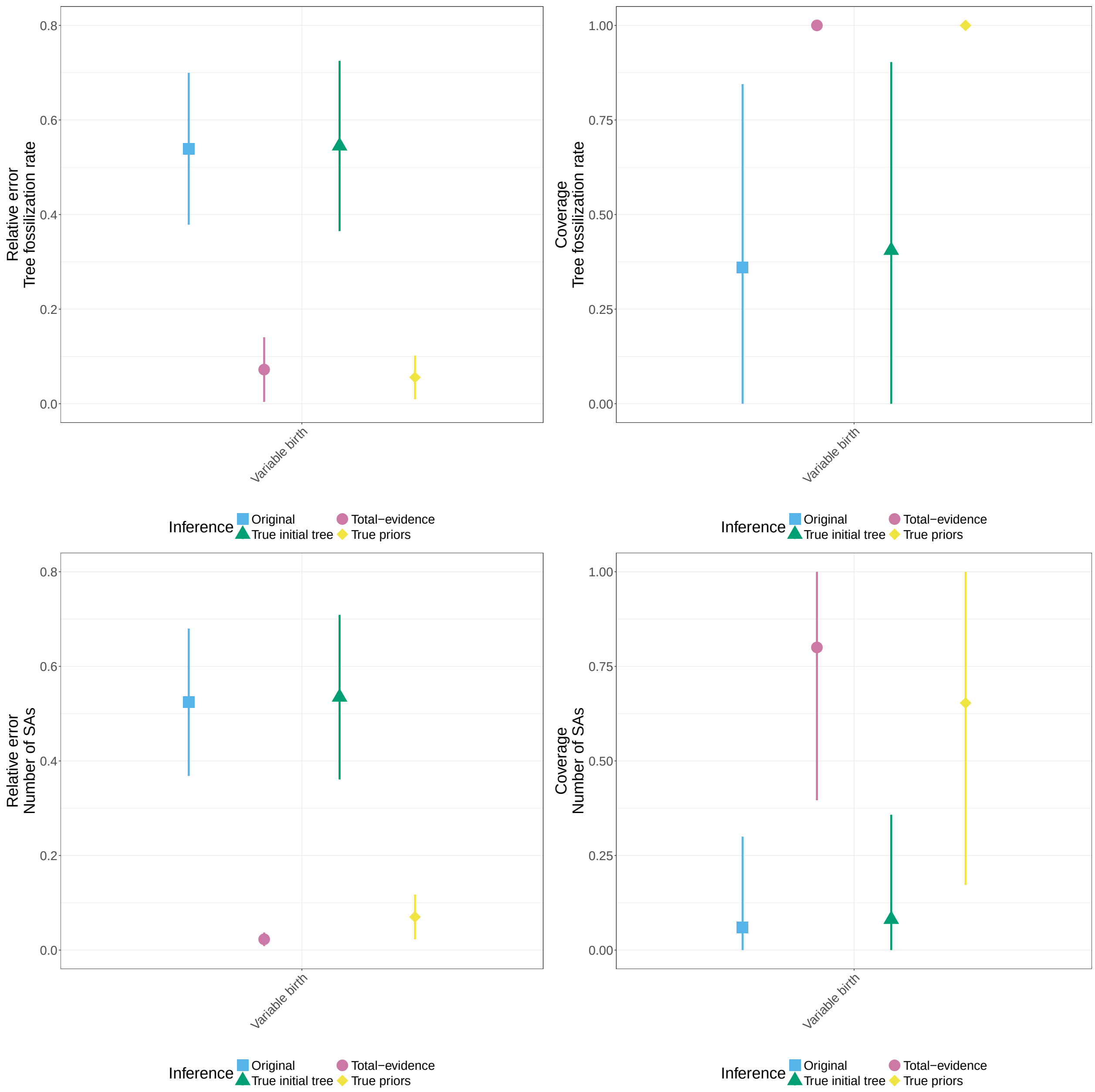


Figure 7: Relative error and 95% HPD coverage of the estimated fossilization rate averaged over the phylogeny (top row) and the estimated number of SAs (bottom row), by inference under the MTFBD process. The inference was run either with topological constraints on the fossil placement (blue, green, yellow) or using a morphological character matrix (pink). The plots show the average and standard deviation over the 50 replicates for the dataset with variable birth rate.

extinct tips. Similar to the results obtained on the full tree, the fossilization rate can be accurately estimated by all types of inference, although coverage is low for the MTFBD inference with topological constraints on all datasets. Coverage is also low for the FBD inferences on datasets which contain variation in the fossilization rates.

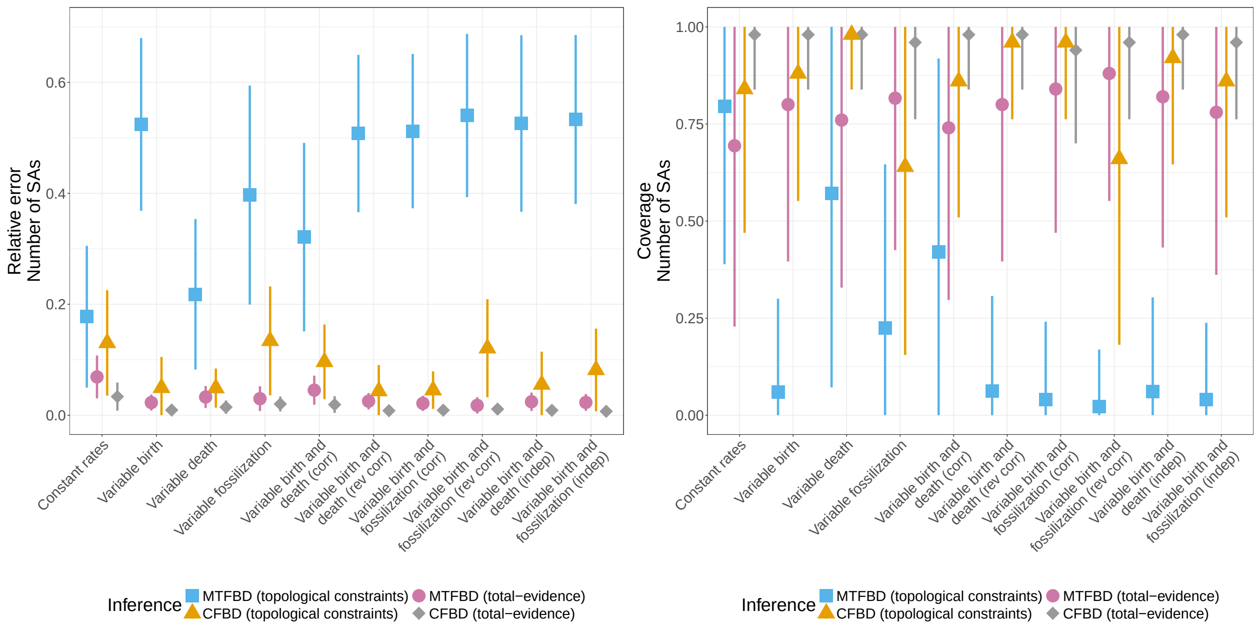


Figure 8: Relative error and 95% HPD coverage of the estimated number of fossils which are sampled ancestors, by inference under a constant-rate FBD process (orange and gray) or the MTFBD process (blue and pink). The plots show the average and standard deviation over the 50 replicates for each dataset.

As in the previous section, results on the tip birth, death and fossilization rate estimates suggest that the inference on specific tips is less reliable than for the global pattern across the entire tree. In particular, when a dataset includes variation on a specific rate, FBD inferences show low coverage values for tip estimates for that rate. This shows that although the FBD inference provides accurate rate estimates on average, it cannot adjust to individual variations between tips, as expected from a constant-rate model. As in the previous section, we note that tip rates can be driven by type changes which are very close to the tips, and thus not identifiable from the phylogeny even when using an MTFBD model (as was previously observed in Barido-Sottani et al. (2020)).

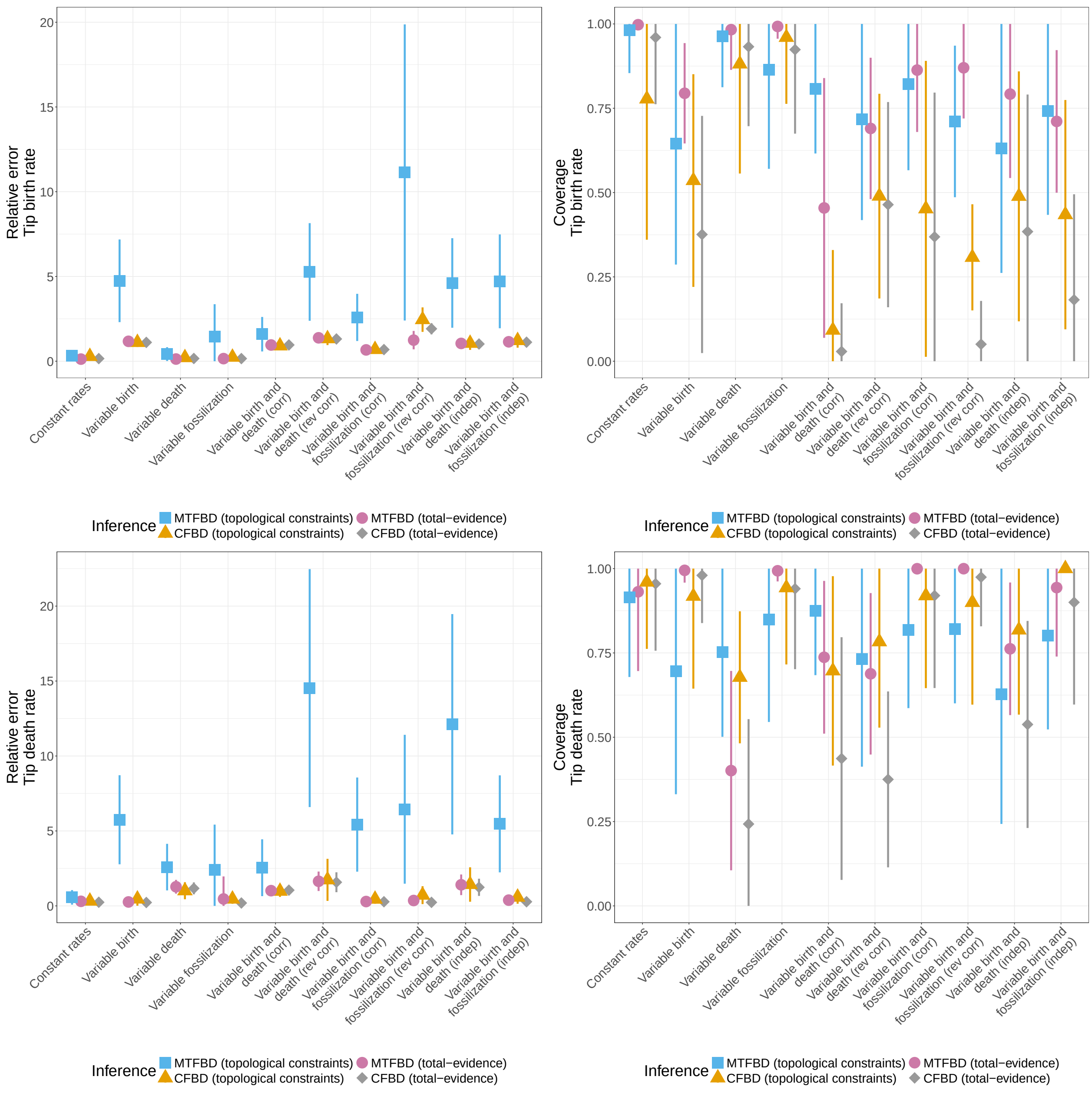


Figure 9: Relative error and 95% HPD coverage of the estimated birth rate (top row) and death rate (bottom row), by inference under a constant-rate FBD process (orange and gray) or the MTFBD process (blue and pink), averaged over all extant and extinct tips. The inference was run either with topological constraints on the fossil placement (orange and blue) or using a morphological character matrix (gray and pink). The plots show the average and standard deviation over the 50 replicates for each dataset.

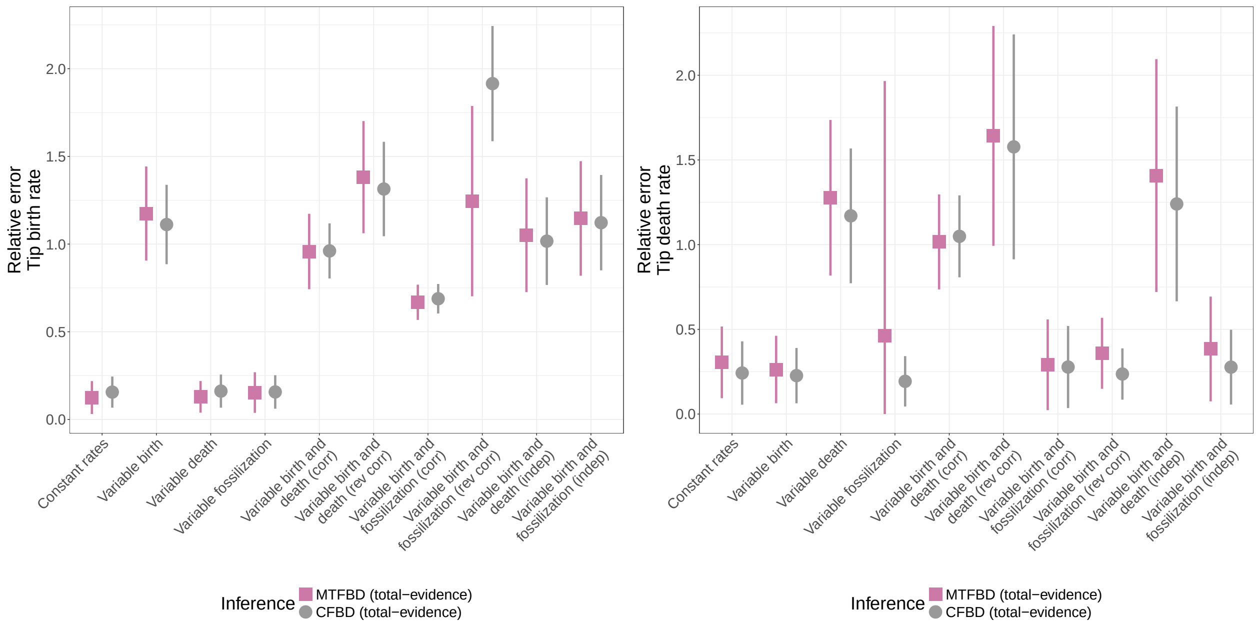


Figure 10: Relative error of the estimated birth rate (left) and death rate (right), by **total evidence** inference under a constant-rate FBD process (gray) or the MTFBD process (pink), averaged over all extant and extinct tips. The plots show the average and standard deviation over the 50 replicates for each dataset.

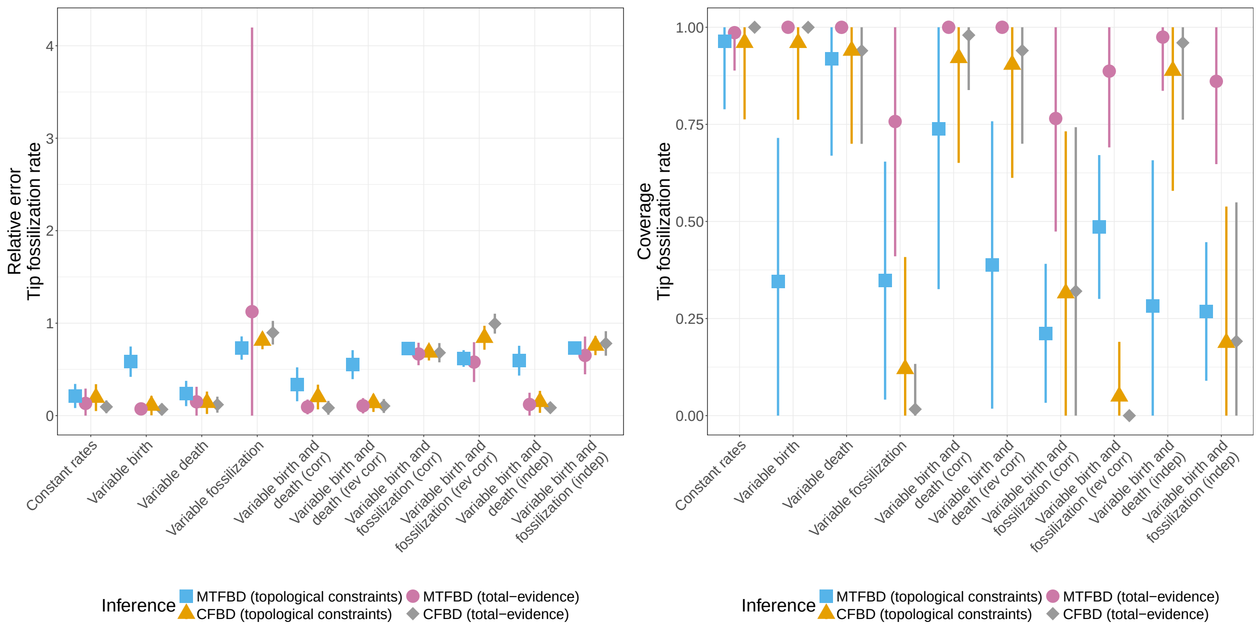


Figure 11: Relative error and 95% HPD coverage of the estimated fossilization rate, by inference under a constant-rate FBD process (orange and gray) or the MTFBD process (blue and pink), averaged over all extant and extinct tips. The inference was run either with topological constraints on the fossil placement (orange and blue) or using a morphological character matrix (gray and pink). The plots show the average and standard deviation over the 50 replicates for each dataset.

#### 3 Empirical dataset

Figure 12 shows the summary tree obtained under the constant-rate FBD process. The median estimated rates for the constant-rate process are 0.56 for the birth rate, 0.54 for the death rate and 0.022 for the fossil sampling rate.

Figure 13 shows the MCC summary tree estimated using the MTFBD model coloured by the mode of the estimated fossil sampling rate per edge.

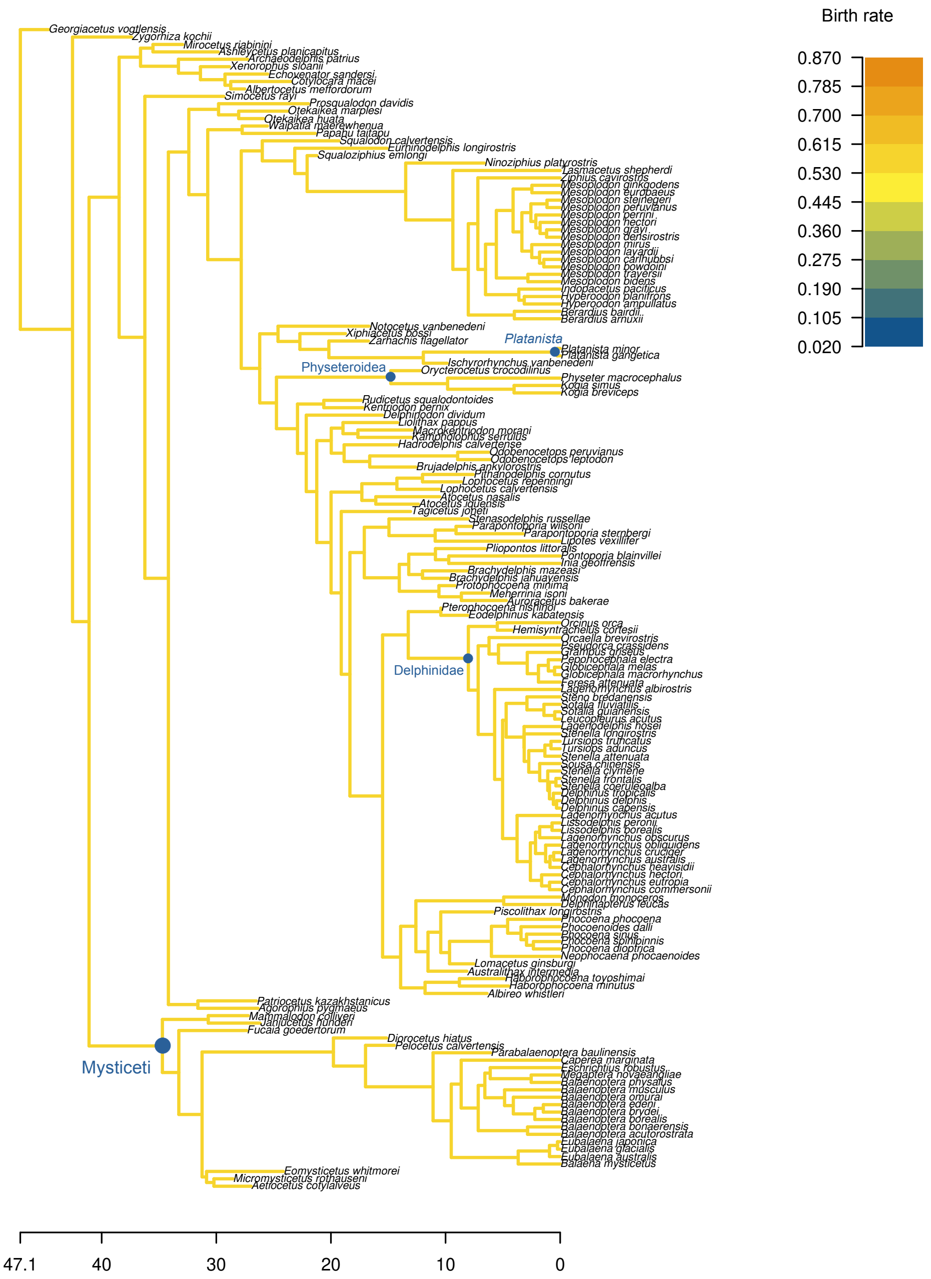


Figure 12: MCC summary tree of the cetaceans phylogeny inferred under a constant-rate FBD process, coloured by the median estimated birth rate (constant over all the phylogeny). The MRCA of clades mentioned in the main text is indicated by blue dots.

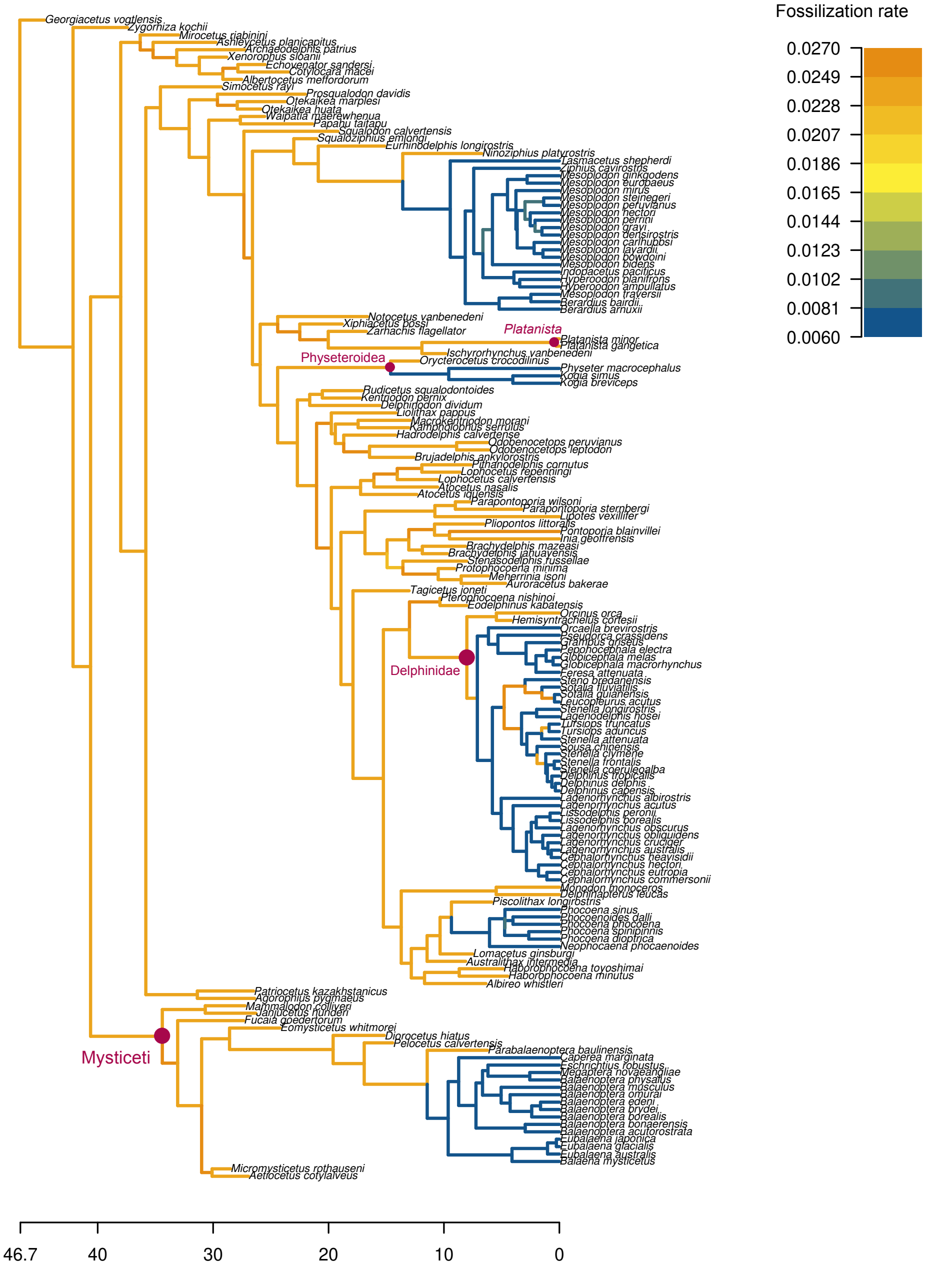


Figure 13: MCC summary tree of the cetaceans phylogeny inferred under a combined MTFBD model. Each edge is coloured by the mode of the estimated fossil sampling rate. The MRCA of clades mentioned in the main text is indicated by pink dots.
